# Evaluation of the Efficacy of the Hypocretin/orexin Receptor Agonists TAK-925 and ARN-776 in Narcoleptic *Orexin/tTA; TetO-DTA* Mice

**DOI:** 10.1101/2022.12.09.519796

**Authors:** Yu Sun, Alok Ranjan, Ryan Tisdale, Shun-Chieh Ma, Sunmee Park, Meghan Haire, Jasmine Heu, Stephen R. Morairty, Xiaoyu Wang, Daniel M. Rosenbaum, Noelle S. Williams, Jef K. De Brabander, Thomas S. Kilduff

## Abstract

The sleep disorder Narcolepsy, a hypocretin deficiency disorder thought to be due to degeneration of hypothalamic hypocretin/orexin neurons, is currently treated symptomatically. We evaluated the efficacy of two small molecule hypocretin/orexin receptor-2 (HCRTR2) agonists in narcoleptic male *orexin/tTA; TetO-DTA* mice. TAK- 925 (1-10 mg/kg, s.c.) and ARN-776 (1-10 mg/kg, i.p.) were injected 15 min before dark onset in a repeated measures design. EEG, EMG, subcutaneous temperature (T_sc_) and activity were recorded by telemetry; recordings for the first 6-h of the dark period were scored for sleep/wake and cataplexy. At all doses tested, TAK-925 and ARN-776 caused continuous wakefulness and eliminated sleep for the first hour. Both TAK-925 and ARN-776 caused dose-related delays in NREM sleep onset. All doses of TAK-925 and all but the lowest dose of ARN-776 eliminated cataplexy during the first hour after treatment; the anti-cataplectic effect of TAK-925 persisted into the 2^nd^ hour for the highest dose. TAK-925 and ARN-776 also reduced the cumulative amount of cataplexy during the 6-h post-dosing period. The acute increase in wakefulness produced by both HCRTR2 agonists was characterized by increased spectral power in the gamma EEG band. Although neither compound provoked a NREM sleep rebound, both compounds affected NREM EEG during the 2^nd^ hour post-dosing. TAK-925 and ARN-776 also increased gross motor activity, running wheel activity and T_sc_, suggesting that the wake-promoting and sleep-suppressing activities of these compounds could be a consequence of hyperactivity. Nonetheless, the anti-cataplectic activity of TAK-925 and ARN-776 is encouraging for the development of HcrtR2 agonists.

## Introduction

The sleep disorder narcolepsy afflicts 1 in 2,000 people and is characterized by excessive daytime sleepiness (EDS), abnormalities of rapid-eye-movement (REM) sleep, and cataplexy (the sudden loss of muscle tone triggered by emotions). The hypocretin/orexin peptides are synthesized in the perifornical and posterior lateral regions of the human hypothalamus. The two peptides, alternatively called hypocretin-1 or orexin-A and hypocretin-2 or orexin-B, have differential affinities for two receptors, HCRTR1 or OX1R and HCRTR2 or OX2R. Cerebrospinal fluid Hcrt1/orexin-A levels are reduced in people with NT1 relative to controls [1–3]. Degeneration of the hypocretin/orexin (Hcrt) neurons [4, 5] by an immune-related mechanism [6–8] is currently thought to be the proximate cause of narcolepsy type 1 (NT1) but epigenetic silencing of Hcrt expression has recently been suggested [9]. Regardless of the cause, these results suggest NT1 is a disease caused by neuropeptide deficiency.

In animal models, mutations in *Hcrtr2* [10], knockout of the Hcrt precursor protein [11], Hcrt receptors [12–15], or genetic ablation of Hcrt neurons [16–19] all result in a narcoleptic phenotype including cataplexy, sleep/wake fragmentation and increased REM sleep propensity, further linking disrupted Hcrt signaling and narcolepsy. Genetic and pharmacological rescue experiments [20–25] have established the proof-of-concept for replacement of Hcrt neurotransmission as the ideal therapy for narcolepsy. Expression of HCRTR2, the pharmacological target for such therapy, appears intact in humans with narcolepsy [26].

Intracerebroventricular and intrathecal administration of HCRT1/orexin-A increases wake and suppresses cataplexy in hypocretin/orexin neuron-ablated and hypocretin/orexin peptide-deficient mouse models of narcolepsy [20, 27] but not in HCRTR2/OX2R-deficient narcoleptic canines [28], indicating the potential for selective HCRTR2/OX2R agonists in the treatment of narcolepsy and other disorders of excessive sleepiness. Systemic administration of the selective HCRTR2/OX2R agonist, YNT-185 [29], reduced sleep onset REM periods (SOREMs) in hypocretin/orexin peptide-deficient and hypocretin/orexin neuron-ablated mice but not in hypocretin/orexin receptor-deficient mice [30]. These results provided the proof-of-concept for hypocretin/orexin replacement therapy with HCRTR2/OX2R agonists for NT1.

Peripheral administration of YNT185 also promoted wakefulness in hypocretin/orexin-deficient, hypocretin/orexin neuron-ablated and wild-type mice, suggesting that hypocretin/orexin receptor agonists may also be useful for treating sleepiness due to NT1 and other causes. However, YNT185 has limited *in vivo* efficacy and thus appears unsuitable for further clinical development.

Takeda described the HCRTR2/OX2R-selective agonist TAK-925 with an EC_50_ on OX2R = 5.5 nM and >5,000-fold selectivity over HCRTR1/OX1R [31]. When injected subcutaneously, TAK-925 had wake-promoting effects in wild-type and OX2R knockout (KO) mice, a strain that exhibits narcoleptic symptomatology [12]. TAK-925 has since been assessed in human trials (https://clinicaltrials.gov/ct2/show/NCT05180890) but requires intravenous administration. Curiously, although TAK-925 appears to be an effective OX2R agonist and OX2R KO mice are a mouse model of narcolepsy [32], the Takeda paper [31] did not address the efficacy of TAK-925 on cataplexy, the pathognomonic symptom of this disorder. This paper also left several questions unanswered, specifically, (1) why was the sleep/wake data reported for only 1 hr post-injection? (2) Given that TAK-925 is wake-promoting, what effects does it have on locomotor activity? (3) What effects does TAK-925 have on other aspects of physiology such as body temperature? To address these questions, we undertook a study of the efficacy of TAK-925 in a validated mouse model of NT1 [18, 19, 33, 34]. In addition, we sought to determine whether a related analog, ARN-776 [35], was superior to TAK-925 in any of these physiological parameters and assessed whether this analog might avoid the necessity for subcutaneous administration in mouse studies [31] and intravenous administration of TAK-925 in human trials. Because cataplexy is the pathognomonic symptom of NT1 and cataplexy is much more prevalent in the dark period when the mice are active than in the light period, we adminstered these compounds just prior to light offset. We find that both TAK-925 and ARN-776 acutely increase wakefulness and reduce cataplexy but also increase activity and subcutaneous temperature (T_sc_) during the first post-dosing hour, suggesting that the wake-promoting effects are a consequence of increased activity.

## Methods

All experimental procedures were approved by the Institutional Animal Care and Use Committee at SRI International or UT Southwestern Medical Center and were conducted in accordance with the principles set forth in the *Guide for Care and Use of Laboratory Animals*.

### Animals

CD1 mice used for pharmacokinetic experiments were from Charles River Laboratories. Male “DTA mice” used for EEG/EMG experiments were the double transgenic offspring of *orexin/tTA* mice (C57BL/6-Tg(*hOX-tTA*)1Ahky), which express the tetracycline transactivator (tTA) exclusively in Hcrt neurons [19], and B6.Cg-Tg(*tetO-DTA*)1Gfi/J mice (JAX #008468), which express a diphtheria toxin A (DTA) fragment in the absence of dietary doxycycline. Both parental strains were from a C57BL/6J genetic background. Parental strains and offspring used for EEG/EMG recording were maintained on a diet (Envigo T-7012, 200 Doxycycline) containing doxycycline (DOX(+) condition) to repress transgene expression until neurodegeneration was desired. All mice were maintained on a LD12:12 light:dark cycle with food and water *ad libitum*; mice used for EEG/EMG recordings had access to running wheels in their home cage.

### Surgical Procedures

Male DTA mice (N = 7) were aged 11±1 weeks (23±2 g) at the time of surgery. Mice were anesthetized with isoflurane and sterile telemetry transmitters (HD-X02, DSI, St Paul, MN) were placed subcutaneously. Biopotential leads were routed subcutaneously to the head, and both EMG leads were positioned through the right nuchal muscles. Cranial holes were drilled through the skull at -2.0 mm AP from bregma and 2.0 mm ML and on the midline at -1 mm AP from lambda.

The two biopotential leads used as EEG electrodes were inserted into these holes and affixed to the skull with dental acrylic. The incision was closed with absorbable suture. Analgesia was managed with meloxicam (5 mg/kg, s.c.) and buprenorphine (0.05 mg/kg, s.c.) upon emergence from anesthesia and for the first day post-surgery.

Meloxicam (5 mg/kg, s.c., q.d.) was continued for 2 d post-surgery. Mice were monitored daily for 14 days post-surgery and any remaining suture material was removed at the end of this monitoring period. Three weeks after surgery, DTA mice were switched to normal chow (Dox(-) condition) to induce expression of the DTA transgene specifically in the Hcrt neurons and thereby initiate degeneration of these cells [19]. After a 42 d (6 wk) degeneration period, Dox(+) chow was reintroduced to minimize further degeneration and the experimental treatments began.

### Drugs

Methyl (2R,3S)-3-[(methylsulfonyl)amino]-2- {[(cis-4- phenylcyclohexyl)oxy]methyl}piperidine-1-carboxylate (TAK-925; Fig.1a) and (2R,3S)-N-ethyl-2- {[(cis-4-isopropylcyclohexyl)oxy]methyl}-3-[(methylsulfonyl)amino]piperidine-1- carboxamide (ARN-776; Fig. 1b) were synthesized at UT Southwestern Medical Center according to published procedures [35] (see Supplementary Material). These compounds correspond to structures 5 and 2, respectively, from US Patent US 2017/0226137 A1 [35]. Based on results of the pharmacokinetic experiments described below, TAK-925 was dissolved in 10% DMSO / 90% 0.5% methycellulose (“TAK-925 vehicle”) and delivered s.c. and ARN-776 was dissolved in 10% DMSO / 10% Kolliphor EL / 20% PEG400 / 60% dH_2_O (“ARN-776 vehicle”) and delivered i.p. for the EEG/EMG studies. All solutions were prepared fresh each experimental day and administered at 10 mL/kg final volume.

**Fig. 1.**
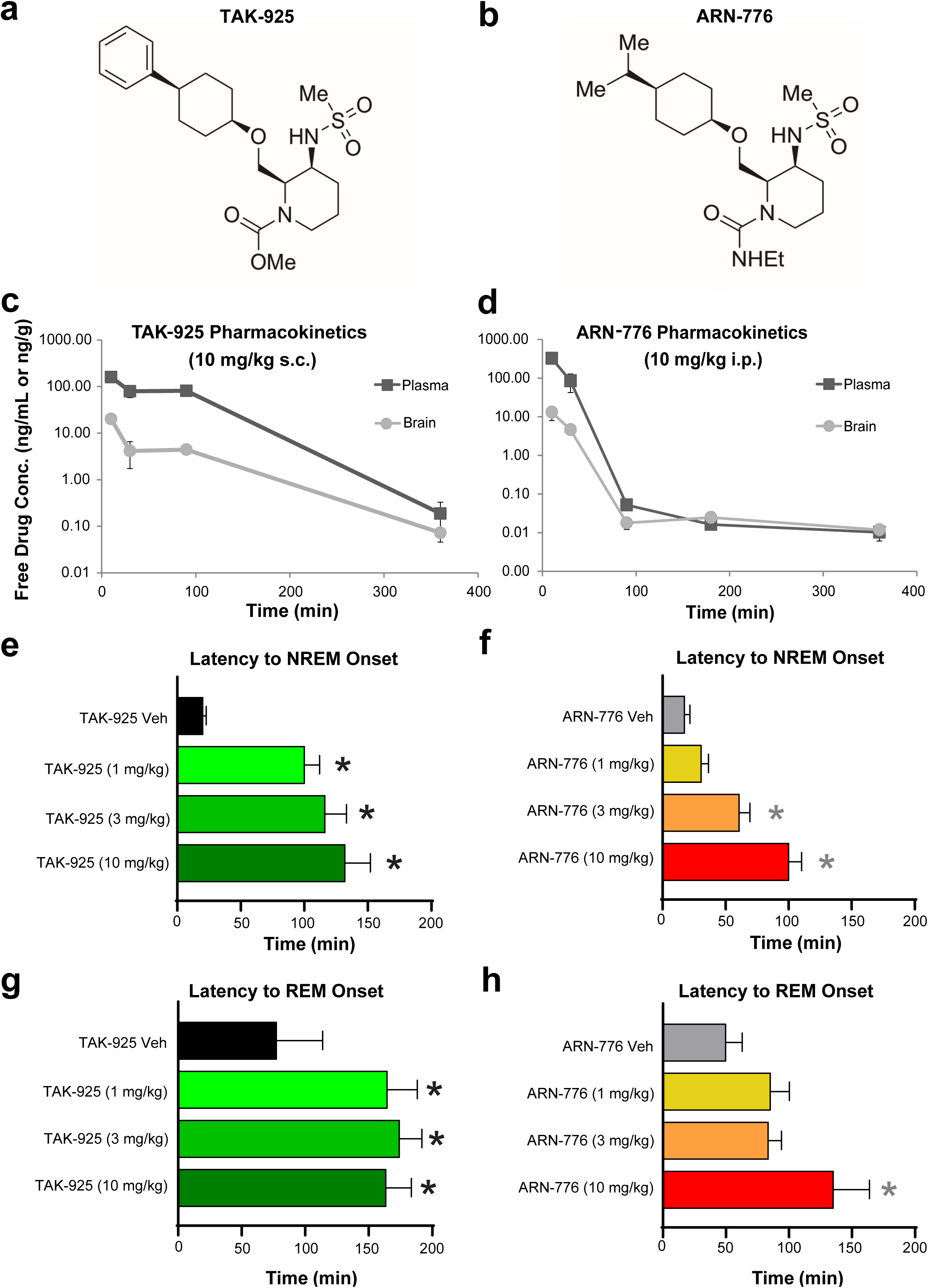
Chemical structures of (**a**) TAK-925 and (**b**) ARN-776. (**c**) Pharmacokinetics of free TAK-925 in CD-1 mice when dosed at 10 mg/kg, s.c. **(d)** Free ARN-776 pharmacokinetics when CD-1 mice were dosed at 10 mg/kg, i.p. (**e**) Latency to NREM sleep in narcoleptic male *orexin/tTA; TetO-DTA* mice after treatment with TAK-925 and **(f)** ARN-776. (**g**) Latency to REM sleep after treatment with TAK-925 and **(h)** ARN-776. Values are mean ± SEM. * *p* < 0.05 compared to Vehicle by 1-way ANOVA followed by the Holm-Sidak *post hoc* test. Abbreviations: Veh, vehicle.

### Pharmacokinetic experiments

CD1 mice were dosed with TAK-925 (10 mg/kg, s.c.) or with ARN-776 (10 mg/kg, i.p.) formulated as described above. At the post-dose times indicated in Fig. 1c and 1d, mice were euthanized with an inhalation overdose of CO_2_ and blood and brain were harvested. Plasma was isolated from blood samples that had been collected using acidified citrate dextrose anticoagulant, and brains were snap frozen after weighing. Brain tissue was homogenized in PBS and both plasma and brain were subject to a simple precipitation step with a 2x volume of methanol to precipitate protein prior to analysis by LC-MS/MS using a fit-for-purpose multiple reaction monitoring method on a Sciex 4000 or 3200 QTRAP™ coupled to a Shimadzu Prominence LC. A simple gradient in methanol/water with 0.1% formic acid (TAK-925 buffers also included 2 mM NH_4_ acetate) using an Agilent C18 XDB column (5 micron, 50 x 4.6 mm) was employed. N-benzylbenzamide was used as an internal standard and concentrations were determined in comparison to standard curves made by spiking compound into blank plasma or brain homogenate. The limit of quantitation was 0.5 ng/mL for TAK-925 and 0.05 ng/mL for ARN-776. To determine free drug levels, the fraction of unbound drug (fu) was determined using rapid equilibrium dialysis (RED). Commercial CD1 mouse plasma (Bioreclamation) or blank brain homogenate derived in house from CD1 mice were both diluted 20x in PBS, pH adjusted to 7.4, and incubated with compound at 5 µM in a Thermo Scientific Pierce RED device single use plate with inserts on an orbital shaker at 37°C, 5% CO_2_, 75% relative humidity for 6-h. Upon harvest, samples from each side of the dialysis chamber were matrix-matched and processed for analysis as described above for PK samples. Percent Protein Binding (%PPB) was determined according to the following equation:

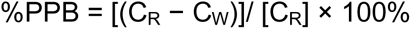

where C_R_ is total drug concentration in plasma in red chamber and C_W_ is free drug concentration in white chamber. All concentrations are approximated by peak area ratio of analytes/IS determined by LC-MS/MS. A correction [37] was applied for matrix dilution as follows:

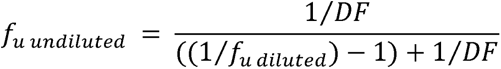

where

DF – dilution factor for matrix dilution

f_u undiluted_ – fraction unbound after correcting for matrix dilution

f_u diluted_ – fraction unbound of diluted matrix

Total plasma and brain concentrations were multiplied by the determined free fraction in each matrix before plotting. Pharmacokinetic parameters were determined using noncompartmental analysis with sparse sampling using WinNonlin (Pharsight).

### Drug treatments

We utilized a repeated measures design in which all animals received all treatments. Although DTA mice received 9 treatments in balanced order over a 4 week period with at least 3 d between dosings, only 8 treatments were analyzed for this study: TAK-925 vehicle, 1, 3 and 10 mg/kg, s.c. and ARN-776 vehicle 1, 3 and 10 mg/kg, i.p. Because cataplexy is the pathognomonic symptom of NT1 and cataplexy is much more prevalent in the dark period, all dosings started 15 min before light offset at ZT12 (ZT0 = 05:00; ZT12 = 17:00).

### EEG, EMG, T_sc_ and activity recording and analysis

Prior to data collection, DTA mice had at least 2 weeks post-surgical recovery and at least 2 weeks adaptation to running wheels, handling and dosing procedures. Throughout the study, mice were housed individually in home cages with access to food, water, nestlets and running wheels *ad libitum*. Room temperature (22 ± 2°C), humidity (50 ± 20% relative humidity), and lighting conditions (LD12:12) were monitored continuously. Animals were inspected daily in accordance with AAALAC and SRI guidelines and the body mass of each mouse was measured weekly. EEG, EMG, T_sc_, and gross motor activity were recorded via telemetry using Ponemah (DSI, St Paul, MN). EEG and EMG were sampled at 500 Hz. Digital videos were recorded at 10 frames per second, 4CIF de-interlacing resolution.

### Data analysis and statistics

EEG/EMG data were classified in 10-s epochs by expert scorers as wakefulness (W), non-Rapid Eye Movement (NREM) sleep, REM sleep, or Cataplexy using NeuroScore (DSI, St. Paul, MN) as in our previous studies [18, 33]. Criteria for Cataplexy were >10 s of EMG atonia, theta-dominated EEG, and video-confirmed behavioral immobility preceded by ≥40 s of wakefulness [38]. Running wheel activity, determined from video recordings, was scored in 10-s epochs. Epochs containing non-physiological signals in the EEG (e.g., signal disconnection, artifacts due to movement, etc) were scored as artifact and excluded from subsequent spectral analysis. “Mixed epochs” containing more than one vigilance state were tagged similarly which also enabled exclusion from the spectral analysis, but such epochs were included in sleep architecture analyses as the majority state during the epoch. The latency to NREM sleep was defined as the time from dose administration until the first 6 consecutive epochs of NREM. The latency to REM sleep was from dose administration until the first 3 consecutive epochs of REM. Data were analyzed as time spent in state per hour and cumulative time spent in each state for 6-h post-dosing. Sleep/wake architecture measures included the duration and the number of bouts for each state. A “bout” of wake, NREM, REM, or Cataplexy was defined as 2 or more consecutive epochs of that state and ended with a single epoch of any other state. For running wheel activity, a bout was defined as 1 or more consecutive epochs ending in a single epoch of any other state.

The EEG power spectrum (0.3-100 Hz) during W, NREM, REM and Cataplexy was analyzed by fast Fourier transform algorithm on all artifact-free epochs. For spectral analyses, a minimum of 6 consecutive epochs of Wake or NREM sleep and a minimum of 3 consecutive epochs of REM sleep was required for inclusion in the analysis. All periods of Cataplexy, regardless of duration, were included in the spectral analysis. A minimum of 3 animals with enough epochs of each state was also required for inclusion in the analysis. EEG spectra were analyzed in 0.122 Hz bins and in standard frequency bands rounded to the nearest Hz (delta: 0.5-4 Hz, theta: 4-9 Hz, alpha: 9-12 Hz, beta: 12-30 Hz, low gamma: 30-60 Hz and high gamma: 60-100 Hz).

For each mouse, power was normalized to the average power per bin during the 6-h vehicle recording. Hourly averages of T_sc_ and activity determined from the transmitters were also analyzed.

Latencies to NREM and REM sleep, REM/NREM ratios, and total time in state were analyzed using 1-way repeated-measures analysis of variance (ANOVA). When 1-way ANOVA indicated significance, the Holm-Sidak *post hoc* test was utilized to compare to the corresponding vehicle (i.e., TAK-925 doses were compared to the TAK-925 vehicle and ARN-776 doses were compared to the ARN-776 vehicle) to identify any doses that were significant. All other data analyses were by 2-way repeated-measures ANOVA with drug treatment and time as factors using data from all eight treatments including both vehicles. When 2-way ANOVA indicated a treatment x time interaction, paired two-tailed *t*-tests (treatment vs. vehicle) were performed to determine the specific hour during which the difference occurred. Statistics were calculated using functions provided in the MATLAB statistics and machine learning toolbox and using GraphPad Prism (ver. 8.4.2). For the EEG spectra data, statistical comparisons were calculated only on the standard frequency bands. The exact *p* values calculated from the *F* distribution are presented whenever possible but, in cases where the *p* values are extremely small, “*p* <” is used.

## Results

### Pharmacokinetic studies

Fig. 1a and 1b provide the chemical structures of TAK-925 and ARN-776, respectively. The results of the pharmacokinetic studies for TAK-925 (10 mg/kg, s.c.) are presented in Fig. 1c and for ARN-776 (10 mg/kg, i.p.) in Fig. 1d. Total drug levels were converted to free drug after multiplication by the free fraction determined for each drug in mouse plasma and brain using rapid equilibrium dialysis. Analysis of these results by WinNonlin revealed brain:blood ratios of 0.07:1 for TAK-925 and 0.04:1 for ARN-776 (Table 1), indicating poor penetration of both molecules across the blood-brain barrier. Regardless, the quantities of TAK-925 and ARN-776 reaching the brain produced significant physiological effects as described below.

**Table 1.** WinNonlin Noncompartmental PK Parameters

| WinNonlin Noncompartmental PK Parameters (free drug<sup>a</sup>) |  |  |  |  |  |
| --- | --- | --- | --- | --- | --- |
| Parameter | Unit | Free TAK-925 (10 mpk SC) |  | Free ARN-776 (10 mpk IP) |  |
|  |  | Plasma | Brain | Plasma | Brain |
| Fraction unbound<sup>b</sup> | N/A | 0.107 | 0.077 | 0.176 | 0.121 |
| Terminal T<sub>1/2</sub> | min | 36 | 53 | 125 | 38 |
| Tmax | min | 10 | 10 | 10 | 10 |
| Cmax | ng/mL or ng/g | 160 | 20 | 326 | 13 |
| AUC<sub>last</sub> | min*ng/mL or min*ng/g | 20,097 | 1433 | 11,482 | 497 |
| Brain:Blood Ratio |  | 0.07:1 |  | 0.04:1 |  |

<sup>a</sup>Total drug concentrations determined by LC-MS/MS were converted to free drug after multiplying by the fraction unbound. PK parameters were determined based on the data presented in Fig. 1C and D using sparse sampling and noncompartmental analysis (NCA) with WinNonlin (Pharsight). Terminal T<sub>1/2</sub> was determined using best fit according to standard parameters provided for NCA in WinNonlin.
<sup>b</sup>Fraction unbound was determined by incubation of 5 µM of each compound in either mouse plasma or brain homogenate using rapid equilibrium dialysis (RED).

### Effects on Sleep Onset

Both TAK-925 and ARN-776 caused dose-related delays in the onset of NREM sleep (*F*_(7, 42)_ *=* 18.67, *p* = 4.86 x 10^-11^). *Post hoc* tests revealed that all three s.c. doses of TAK-925 delayed NREM sleep onset (Fig. 1e) whereas only the 3 mg/kg and 10 mg/kg i.p. doses of ARN-776 delayed NREM (Fig. 1f). ANOVA also indicated that both compounds delayed REM sleep onset (*F*_(7, 42)_ *=* 6.32, *p =* 4.29 x 10^-5^); *post hoc* tests determined that all doses of TAK-925 (Fig. 1g) and the 10 mg/kg dose of ARN-776 (Fig. 1h) delayed REM sleep.

### Hourly Amounts of Sleep/Wake

Two-way ANOVA indicated significant treatment x time effects on the amounts of Wake (*F*_(35, 210)_ = 2.79, *p* = 3.17 x 10^-6^), NREM (*F*_(35, 210)_ = 2.11, *p* = 6.68 x 10^-4^) and REM sleep (*F*_(35, 210)_ = 2.53, *p* = 2.54 x 10^-5^). TAK-925 caused continuous wakefulness (Fig. 2a), eliminated both NREM (Fig. 2d) and REM sleep (Fig. 2g) as well as Cataplexy (Fig. 2j) for the first hour at all doses tested (1-10 mg/kg, s.c.); the reduction of Cataplexy continued into the second post-dosing hour for the 10 mg/kg TAK-925 dose. However, the TAK-925 effects on arousal states were transitory (Fig. 2c, 2f and 2i) and only the highest TAK-925 dose reduced Cataplexy across the entire 6-h recording period (Fig. 2l). The suppression of REM sleep by TAK-925 early in the night was followed by a rebound that reached significance at ZT16, 4 hours post-treatment (Fig. 2g).

**Fig. 2.**
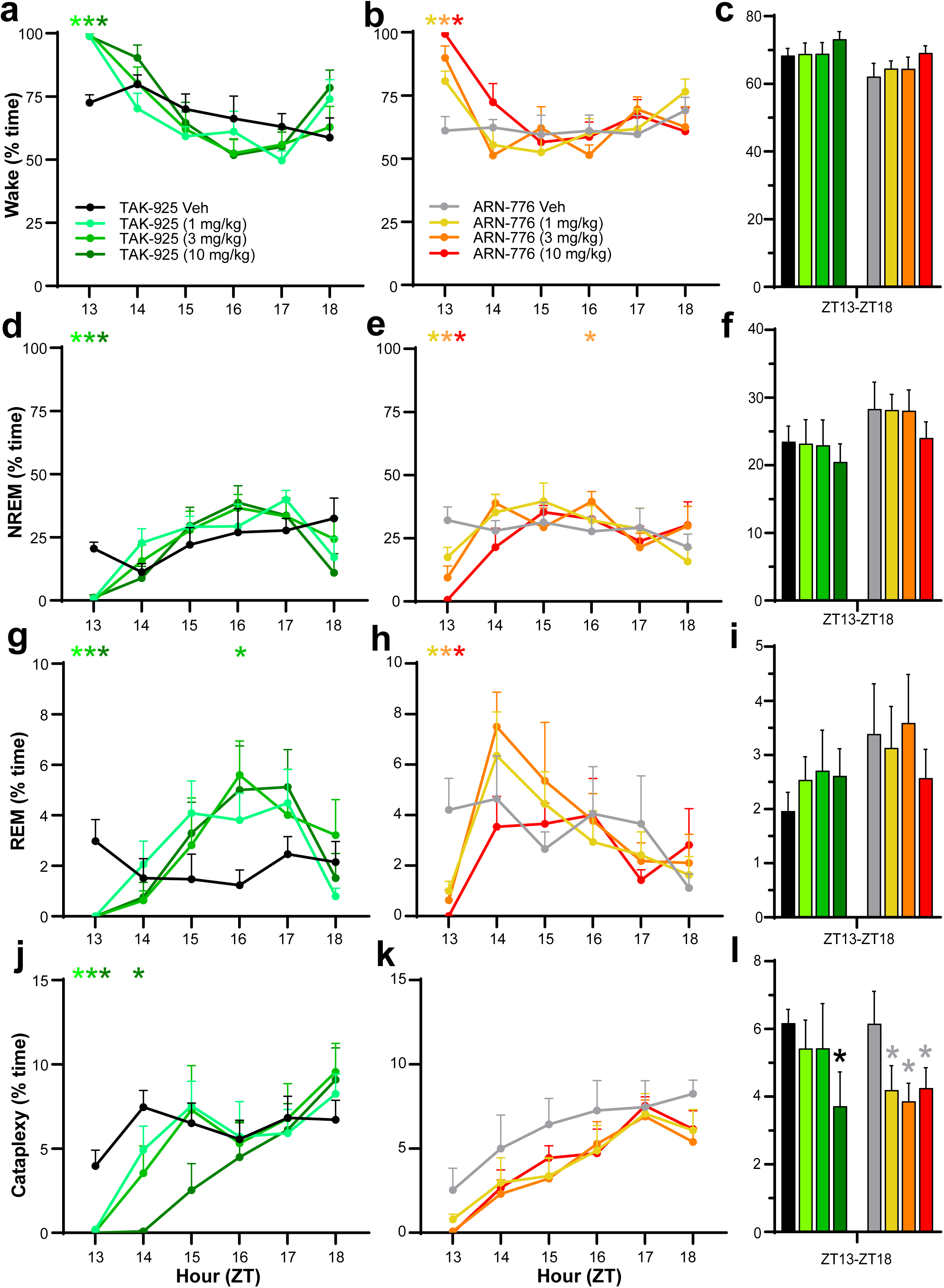
Hourly amounts of Wakefulness (**a, b**), NREM sleep (**d, e**), REM sleep (**g, h**), and cataplexy (**j, k**) in narcoleptic male *orexin/tTA; TetO-DTA* mice treated with TAK- 925 and ARN-776. Percent time in Wake (**c**), NREM sleep (**f**), REM sleep (**i**) and Cataplexy (**l**) summed across the 6-h recording after treatment with TAK-925 and ARN- 776. Values are mean ± SEM. Colored * above hourly graphs indicate significance (*p* < 0.05) during that hour for that dose relative to vehicle treatment. * in **l** denotes significance for that treatment relative to vehicle across the 6-h recording as determined by the Holm-Sidak *post hoc* test. Abbreviations: Veh, vehicle; ZT, Zeitgeber Time.

In contrast to TAK-925, ARN-776 (1-10 mg/kg, i.p.) produced dose-related changes in Wake, NREM and REM sleep during the first hour (Fig. 2b, 2e, 2h) but the reduction in Cataplexy levels did not reach statistical significance (Fig. 2k), likely due to the slightly lower levels of Cataplexy after treatment with the ARN-776 vehicle during that hour than with the TAK-925 vehicle. Across the entire 6-h recording, however, ANOVA revealed a significant treatment effect of ARN-776 on Cataplexy (*F*_(3, 18)_ = 3.928 *p* = 0.0256); the *post hoc* Holm-Sidak multiple comparisons test indicated that all 3 doses of ARN-776 reduced Cataplexy across the 6-h period (Fig. 2l). In further contrast to TAK-925, the suppression of REM sleep by ARN-776 was briefer and the rebound did not reach statistical significance (Fig. 2g vs. 2h).

The REM/NREM ratio was unchanged by TAK-925 or ARN-776 when measured across the entire 6-h recording.

### Cumulative Amounts of Sleep/Wake

#### Treatment effects

The changes described above were reflected in the cumulative amounts of each state (Fig. 3), for which two-way ANOVA revealed significant treatment effects on Wake (*F*_(7, 42)_ = 5.03, *p* = 3.3 x 10^-4^), NREM (*F*_(7, 42)_ = 3.26, *p* = 0.007), REM (*F*_(7, 42)_ = 2.54, *p* = 0.03) and Cataplexy (*F*_(7, 42)_ = 3.27, *p* = 0.007). The initial promotion of Wake (Fig. 3a, 3b) and suppression of both NREM (Fig. 3c, 3d) and REM sleep (Fig. 3e, 3f) by both compounds are readily evident, as is the more sustained effect of the 10 mg/kg dose of ARN-776*. Post hoc* tests found significant effects of the 10 mg/kg ARN-776 dose on Wake (*p* = 0.005) and NREM (*p* = 0.028). Fig. 3g confirms the suppression of Cataplexy by the TAK-925 10 mg/kg dose (*p* = 1.74 x 10^-4^) that is also shown in Fig. 2l and Fig. 3h shows the cumulative suppression of Cataplexy by all 3 doses of ARN-776 that is also apparent in Fig. 2l.

**Fig. 3.**
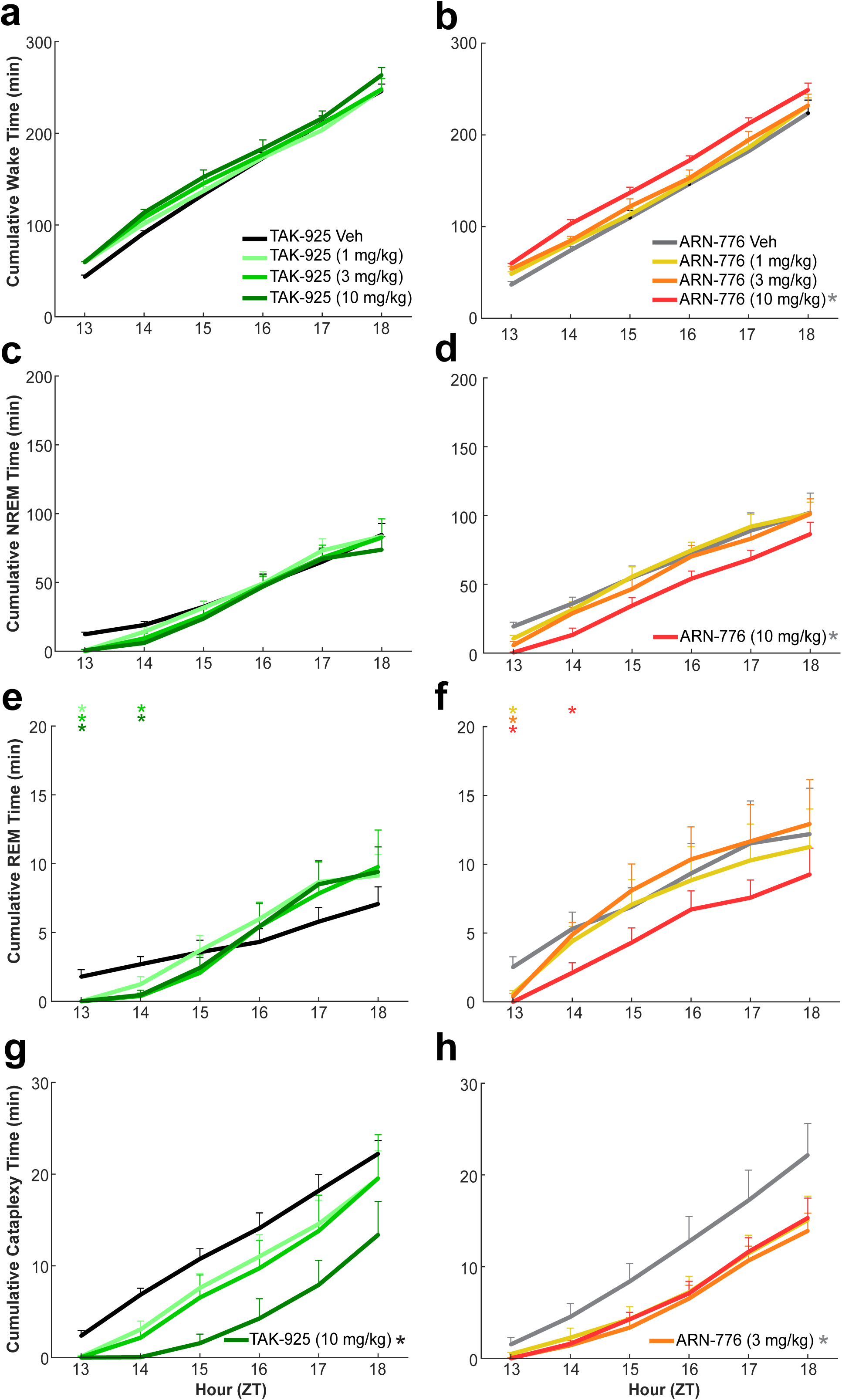
Cumulative time spent in Wakefulness (**a, b**), NREM sleep (**c, d**), REM sleep (**e, f**), and Cataplexy (**g, h**) in male narcoleptic *orexin/tTA; TetO-DTA* mice treated with TAK-925 and ARN-776. Values are mean ± SEM; * in legends denotes significant treatment effect by 2-way ANOVA; colored * above graph indicates significance (*p* < 0.05) during that hour for that dose relative to vehicle treatment. Abbreviations: Veh, vehicle; ZT, Zeitgeber Time.

#### Treatment x Time Effects

Two-way ANOVA revealed a significant treatment x time effect only on the cumulative amount of REM sleep (*F*_(35, 210)_ = 1.61, *p* = 0.02). Fig. 3e clearly illustrates the significant suppression of REM sleep by TAK-925 during the first two hours after dosing that was followed by a later rebound; in contrast, the suppression of REM sleep by ARN-776 at the lower doses was limited to the first post-dosing hour (Fig. 3f).

### Sleep Architecture Measures

#### Bout Durations

The increased wakefulness produced by both Hcrt agonists during the first dosing hour was due to very long Wake bout durations (Fig. 4a, b). Two-way ANOVA confirmed overall treatment effects of TAK-925 and ARN-776 on bout durations for Wake (*F*_(7, 42)_ = 14.48, *p* = 2.08 x 10^-9^), NREM (*F*_(7, 42)_ = 4.85, *p* = 4.52 x 10^-4^), REM (*F*_(7, 42)_ = 4.21, *p* = 1.36 x 10^-3^) and Cataplexy (*F*_(7, 42)_ = 4.12, *p* = 1.58 x 10^-3^). *Post hoc* tests revealed significant treatment effects on Wake bout duration for all doses of both TAK-925 and ARN-776. In contrast, *post hoc* tests did not identify any specific dose of either TAK-925 or ARN-776 that contributed to the overall treatment effects on NREM or REM. The 3 mg/kg dose ARN-776 has a modest effect on Cataplexy bout duration (*p* = 0.046).

**Fig. 4.**
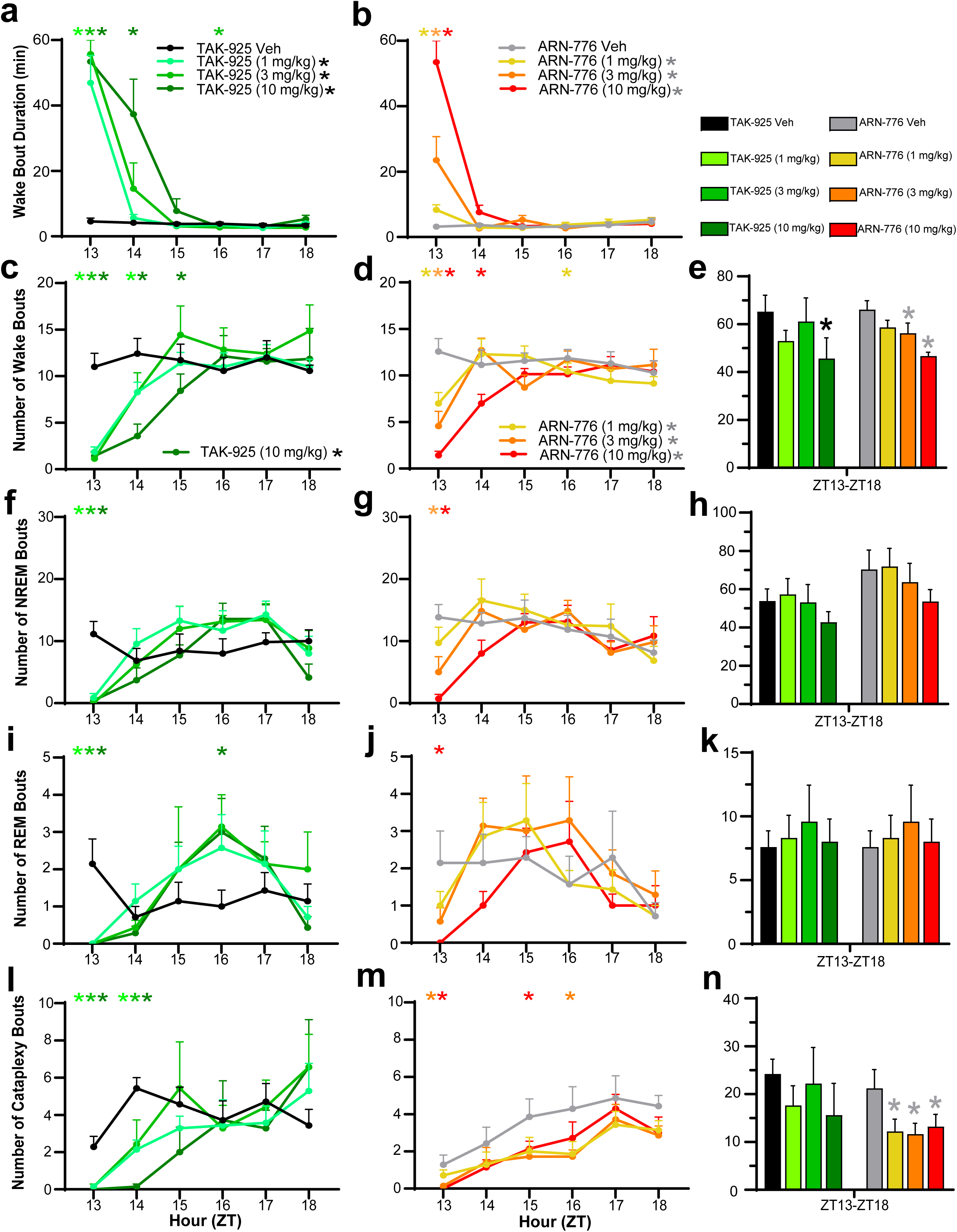
Mean Wake Bout Durations in narcoleptic male *orexin/tTA; TetO-DTA* mice treated with TAK-925 **(a)** and ARN-776 **(b).** Hourly number of bouts of Wakefulness (**c, d**), NREM sleep (**f, g**), REM sleep (**i, j**), and Cataplexy (**l, m**) in narcoleptic male *orexin/tTA; TetO-DTA* mice treated with TAK-925 and ARN-776. Total number of bouts of Wake (**e**), NREM sleep (**h**), REM sleep (**k**) and Cataplexy (**n**) across the 6-h recording after treatment with TAK-925 and ARN-776. Values are mean ± SEM. Colored * above hourly graphs indicate significance (*p* < 0.05) during that hour relative to vehicle treatment. * in legend denotes significance for that treatment across the 6-h recording as determined by 2-way ANOVA. * in **e** and **n** denotes significance for that treatment relative to vehicle across the 6-h recording as determined by *post hoc* Holm-Sidak test. Abbreviations: Veh, vehicle; ZT, Zeitgeber Time.

Two-way ANOVA also revealed a treatment x time interaction on Wake bout duration (*F*_(35, 210)_ = 12.74, *p* < 3.34 x 10^-13^). *Post hoc* tests revealed that all doses of both TAK-925 and ARN-776 increased Wake bout duration during the first post-dosing hour (ZT13), which continued into the second post-dosing hour (ZT14) for the highest dose of TAK-925 (Fig. 4a).

#### Number of Bouts

Two-way ANOVA determined significant treatment effects for TAK-925 and ARN-776 on the number of Wake bouts across the entire 6-h recording (*F_(_*_7, 42)_ = 2.76, *p* = 0.02). The Holm-Sidak *post hoc* tests found that only the TAK-925 10 mg/kg dose reduced the number of Wake bouts across the entire 6-h recording (*p* = 0.013) whereas the 3 mg/kg (*p* = 0.03) and 10 mg/kg (*p* = 0.002) doses of ARN-776 significantly reduced the number of Wake bouts (Fig. 4e).

Two-way ANOVA also revealed significant treatment x time interactions for TAK- 925 and ARN-776 on the number of bouts of Wake (*F*_(35, 210)_ = 4.78, *p* = 3.34 x 10^-13^), NREM (*F*_(35, 210)_ = 2.81, *p* = 2.84 x 10^-6^), REM (*F*_(35, 210)_ = 1.72, *p* = 0.011) and Cataplexy (*F*_(35, 210)_ = 1.80, *p* = 6.32 x 10^-3^). During the first post-dosing hour when wakefulness was continuous, *post hoc* tests revealed that all doses of TAK-925 reduced the number of bouts of Wake (Fig. 4c), NREM (Fig. 4f), REM (Fig. 4i) and Cataplexy (Fig. 4l); the reduction of Wake bouts lasted up to 3h at 10 mg/kg. By contrast, ARN-776 had a dose-related effect on the number of Wake bouts during the first hour but only the 10 mg/kg dose reduced the number of Wake bouts during the second hour (Fig. 4d).

Overall, ARN-776 appeared to have weaker effects than TAK-925 at the doses tested, as only the middle and high doses reduced the number of NREM bouts during the first hour post-dosing (Fig. 4g) and only the 10 mg/kg dose reduced the number of REM bouts (Fig. 4j). Similarly, whereas all doses of TAK-925 reduced the number of Cataplexy bouts for the first two hours (Fig. 4l), only the two highest concentrations of ARN-776 were effective during the first and later hours (Fig. 4m). Nonetheless, the *post hoc* Holm-Sidak test revealed that all doses of ARN-776 (1 mg/kg: *p* = 0.0082; 3 mg/kg: *p* = 0.0078; 10 mg/kg: *p* = 0.0092) reduced the total number of Cataplexy bouts across the entire 6-h period (Fig. 4n).

### EEG Spectra Effects

#### Wakefulness

The acute increase in wakefulness produced by both Hcrt agonists during the first hour post-dosing (ZT13) was characterized by increased spectral power in the alpha and both low and high gamma bands of the EEG (Fig. 5a, 5b and Fig. 6e, 6f, 6i, 6j, 6k and 6l). Two-way ANOVA found significant effects of treatment on the high gamma band (*F*_(7, 42)_ *=* 4.91, *p* = 4.09 x 10^-4^) during Wake across the 6-h recording. *Post hoc* testing found that the 10 mg/kg doses of both TAK-925 (*p* = 0.02, Fig. 6K) and ARN-776 (*p* = 0.04, Fig. 6l) significantly increased high gamma during Wake across the entire recording period.

**Fig. 5.**
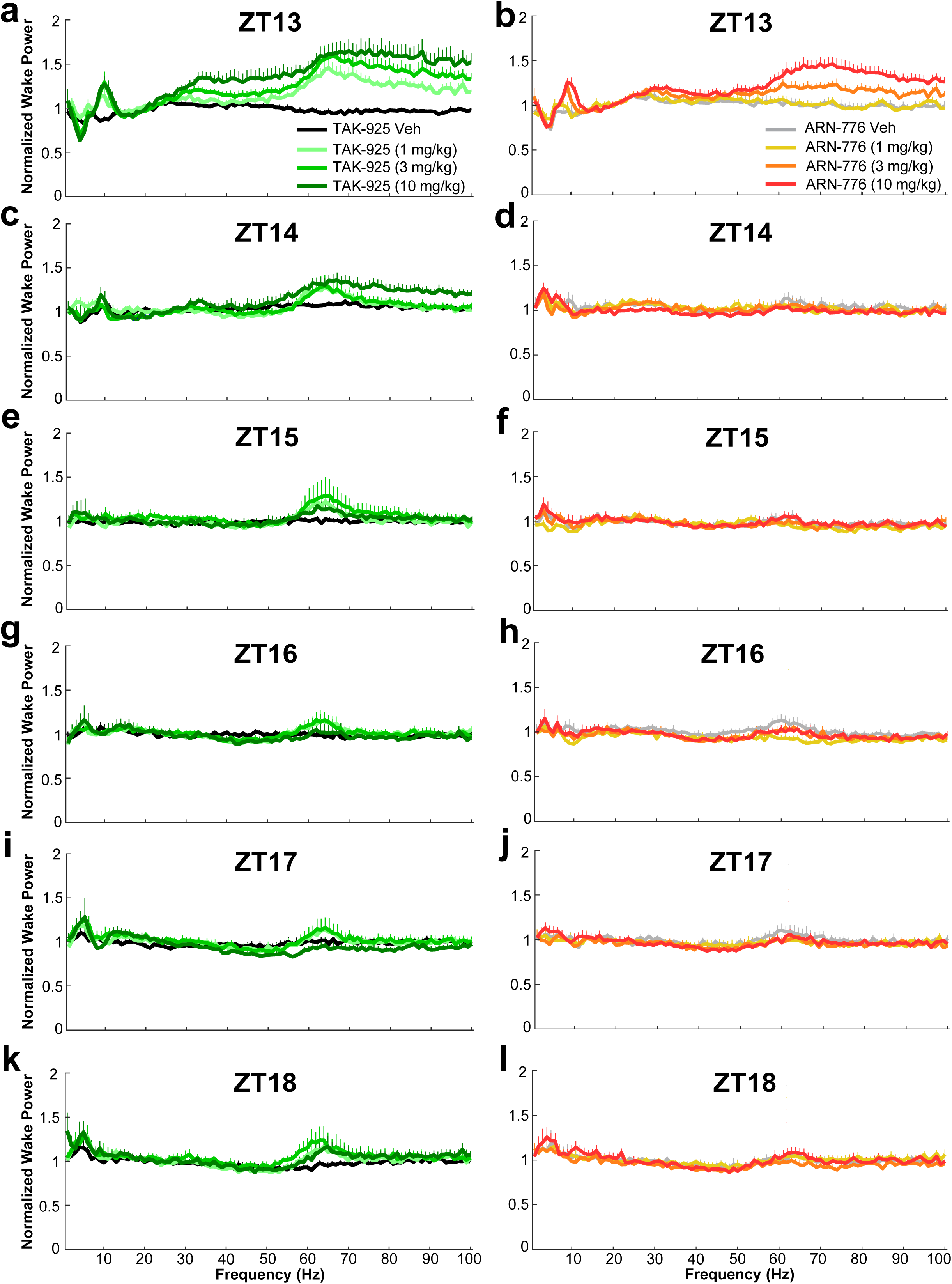
Normalized EEG spectral power (0 – 100 Hz) recorded during wakefulness for the 6-h post-dosing recording period in male narcoleptic *orexin/tTA; TetO-DTA* mice treated with TAK-925 (left column) and ARN-776 (right column). Values are mean ± SEM. Abbreviations: Veh, vehicle; ZT, Zeitgeber Time; Hz, Hertz.

**Fig. 6.**
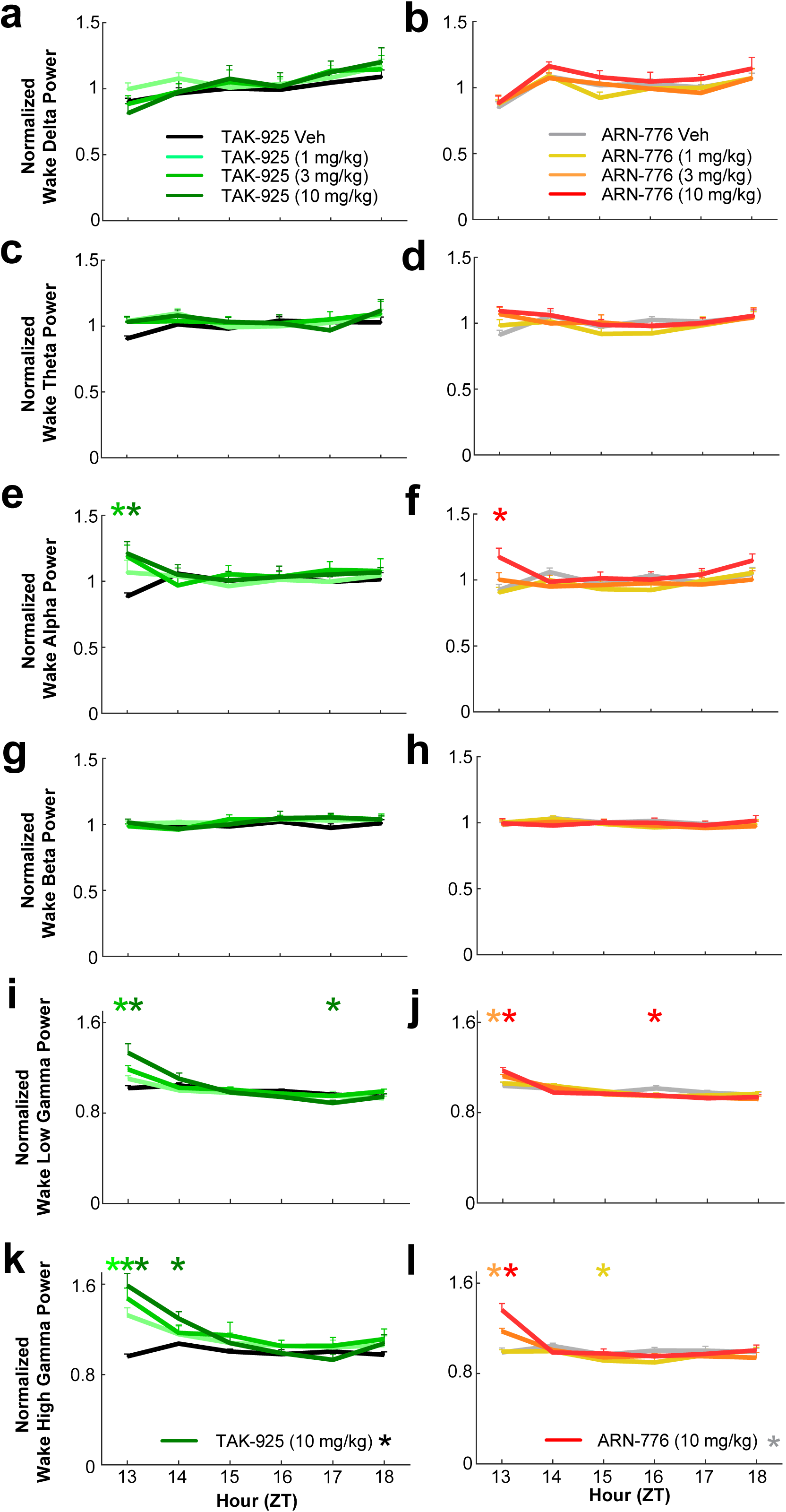
Waking EEG spectra for the conventional bandwidths in narcoleptic male *orexin/tTA; TetO-DTA* mice treated with TAK-925 and ARN-776**. a and b:** delta (0.5-4.0 Hz); **c and d:** theta (4-9 Hz); **e and f:** alpha (9-12 Hz); **g and h:** beta (12-30 Hz); **i and j:** low gamma (30-60 Hz); and **k and l:** high gamma (60-100 Hz). Values are mean ± SEM. Colored * above hourly graphs indicate significance (*p* < 0.05) during that hour relative to vehicle treatment. * in legend denotes significance for that treatment across the 6-h recording as determined by 2-way ANOVA. Abbreviations: Veh, vehicle; ZT, Zeitgeber Time.

ANOVA also revealed significant treatment x time effects on the alpha (*F*_(35, 210)_ *=* 1.99, *p* = 0.0016; Fig. 6e, f), low gamma (*F*_(35, 210)_ *=* 5.07, *p* = 3.41 x 10^-14^; Fig. 6i, j) and high gamma *F*_(35, 210)_ *=* 7.03, *p* < 3.34 x 10^-13^; Fig. 6k, l) ranges of the waking EEG*. Post hoc* tests indicated that the 3 and 10 mg/kg doses of TAK-925 significantly increased spectral power in all three bandwidths during ZT13; the 1 mg/kg dose also significantly increased high gamma power during that hour. The effects of the 10 mg/kg dose continued into the second post-dosing hour (Fig. 6k). *Post hoc* tests also revealed that the 10 mg/kg dose of ARN-776 significantly increased power in all three of these bands during the first post-dosing hour; the 3 mg/kg dose also significantly increased both low (Fig. 6j) and high (Fig. 6l) gamma during that hour (ZT13).

#### NREM Sleep

Across the 6-h recording period, ANOVA revealed significant treatment effects on the delta (*F*_(7, 42)_ = 2.28, *p* = 0.046) and high gamma (*F*_(7, 42)_ = 2.88, *p* = 0.015) bands during NREM sleep (Figs. 7, 8). Unlike Wake, however, *post hoc* testing did not reveal any particular dose of either TAK-925 or ARN-776 that significantly affected these bandwidths.

**Fig. 7.**
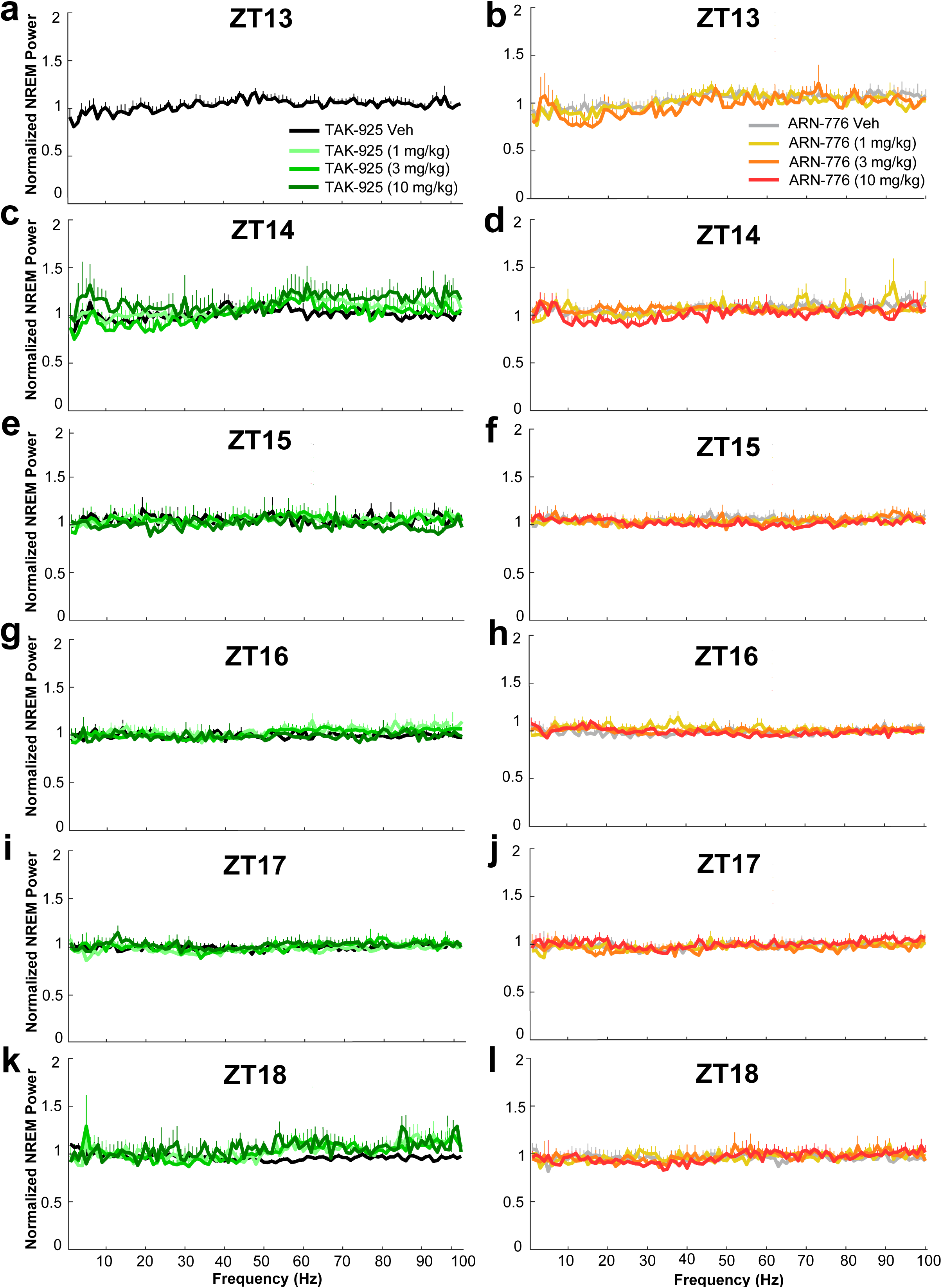
Normalized EEG spectral power (0 – 100 Hz) recorded during NREM sleep for the 6-h post-dosing recording period in male narcoleptic *orexin/tTA; TetO-DTA* mice treated with TAK-925 (left column) and ARN-776 (right column). Values are mean ± SEM. Abbreviations: Veh, vehicle; ZT, Zeitgeber Time; Hz, Hertz.

**Fig. 8.**
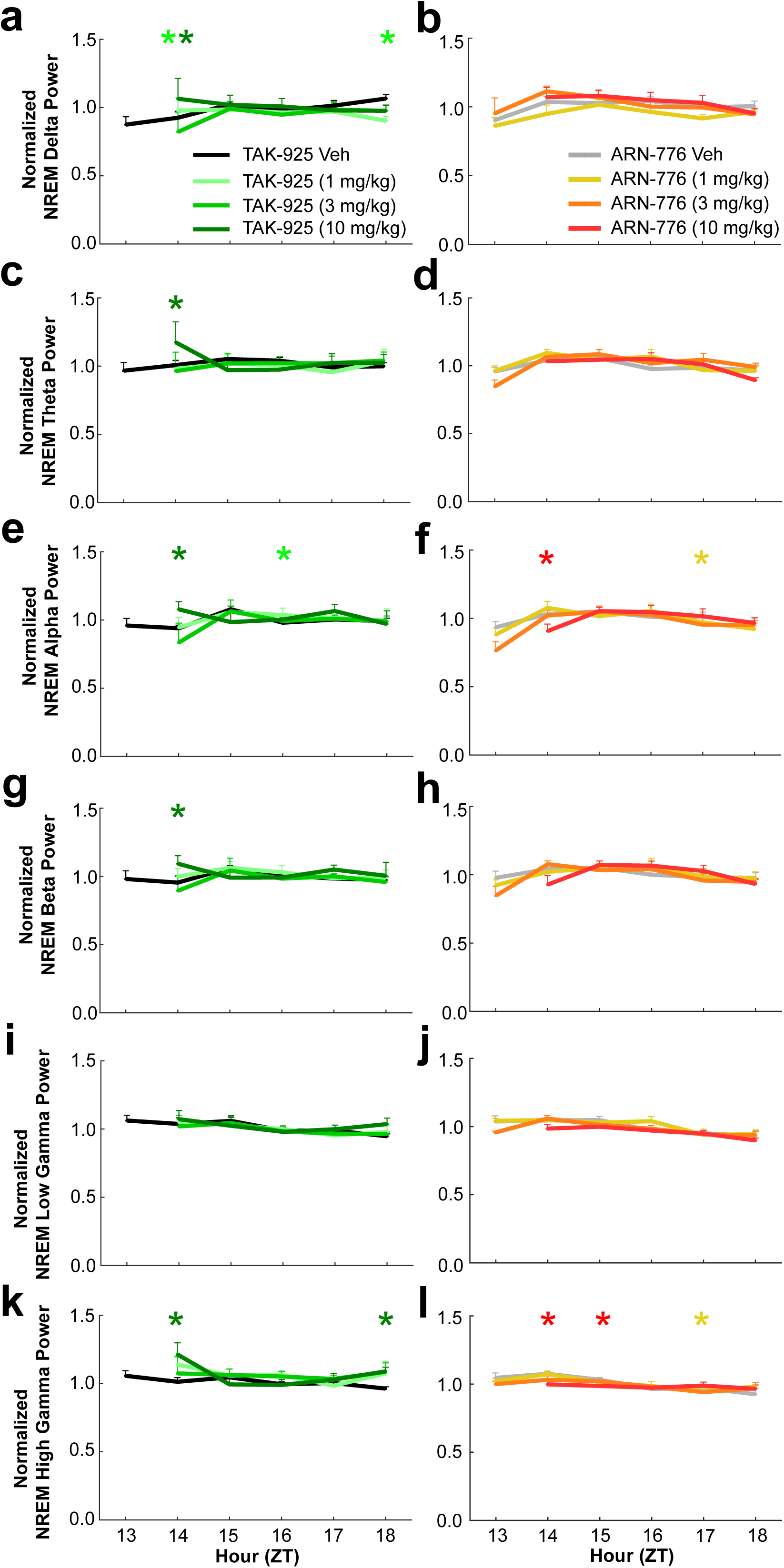
EEG spectra during NREM sleep for the conventional bandwidths in narcoleptic male *orexin/tTA; TetO-DTA* mice treated with TAK-925 and ARN-776**. a and b:** delta (0.5-4.0 Hz); **c and d:** theta (4-9 Hz); **e and f:** alpha (9-12 Hz); **g and h:** beta (12-30 Hz); **i and j:** low gamma (30-60 Hz); and **k and l:** high gamma (60-100 Hz). Values are mean ± SEM. Colored * above hourly graphs indicate significance (*p* < 0.05) during that hour relative to vehicle treatment. Abbreviations: Veh, vehicle; ZT, Zeitgeber Time.

Two-way ANOVA revealed significant treatment x time effects on the delta (*F*_(35, 210)_ = 1.91, *p* = 0.003; Fig. 8a), theta (*F*_(35, 210)_ = 2.31, *p* = 1.43 x 10^-4^; Fig. 8c), alpha (*F*_(35,210)_ = 3.00, *p* = 5.99 x 10^-7^; Fig. 8e, 8f), beta (*F*_(35, 210)_ = 2.08, *p* = 8.63 x 10^-4^; Fig. 8g), and high gamma (*F*_(35, 210)_ = 2.37, *p* = 9.38 x 10^-5^; Fig. 8k, 8l) ranges of the EEG during NREM sleep. *Post hoc* tests indicated that most of the significant effects of TAK-925 on NREM sleep occurred during ZT14, the hour after the complete suppression of sleep by TAK-925. The 10 mg/kg dose of TAK-925 increased spectral power during ZT14 for all five of the EEG bands mentioned above; the 1 mg/kg dose also increased NREM delta during this hour (Fig. 8a). *Post hoc* tests indicated significant reductions in NREM alpha (Fig. 8f) and high gamma (Fig. 8l) during ZT14 after the 10 mg/kg ARN-776 dose; the reduction of high gamma continued into ZT15.

#### REM Sleep

The low levels of REM sleep and the requirement for 3 consecutive epochs of REM for spectral analysis precluded meaningful analysis of the EEG spectra during this state.

#### Cataplexy

Both drugs had strong treatment effects on all EEG bandwidths during cataplexy (Figs. 9, 10) across the 6-h recording period: delta (*F*_(7, 42)_ *=* 7.18, *p* = 1.19 x 10^-5^), theta (*F*_(7, 42)_ *=* 66.26, *p* < 2.62 x 10^-12^), alpha (*F*_(7, 42)_ *=* 21.94, *p* = 3.90 x 10^-12^), beta (*F*_(7, 42)_ *=* 47.43, *p* < 2.62 x 10^-12^), low gamma *F*_(7, 42)_ *=* 132.64, *p* < 2.62 x 10^-12^) and high gamma (*F*_(7, 42)_ = 48.38, *p* < 2.62 x 10^-12^; Figs. 9 and 10). *Post hoc* tests found that the 3 mg/kg dose of TAK-925 reduced theta (*p* = 0.012; Fig. 10c) and that the 3 mg/kg dose of ARN-776 affected high gamma (*p* = 0.004; Fig. 10l) during Cataplexy.

**Fig. 9.**
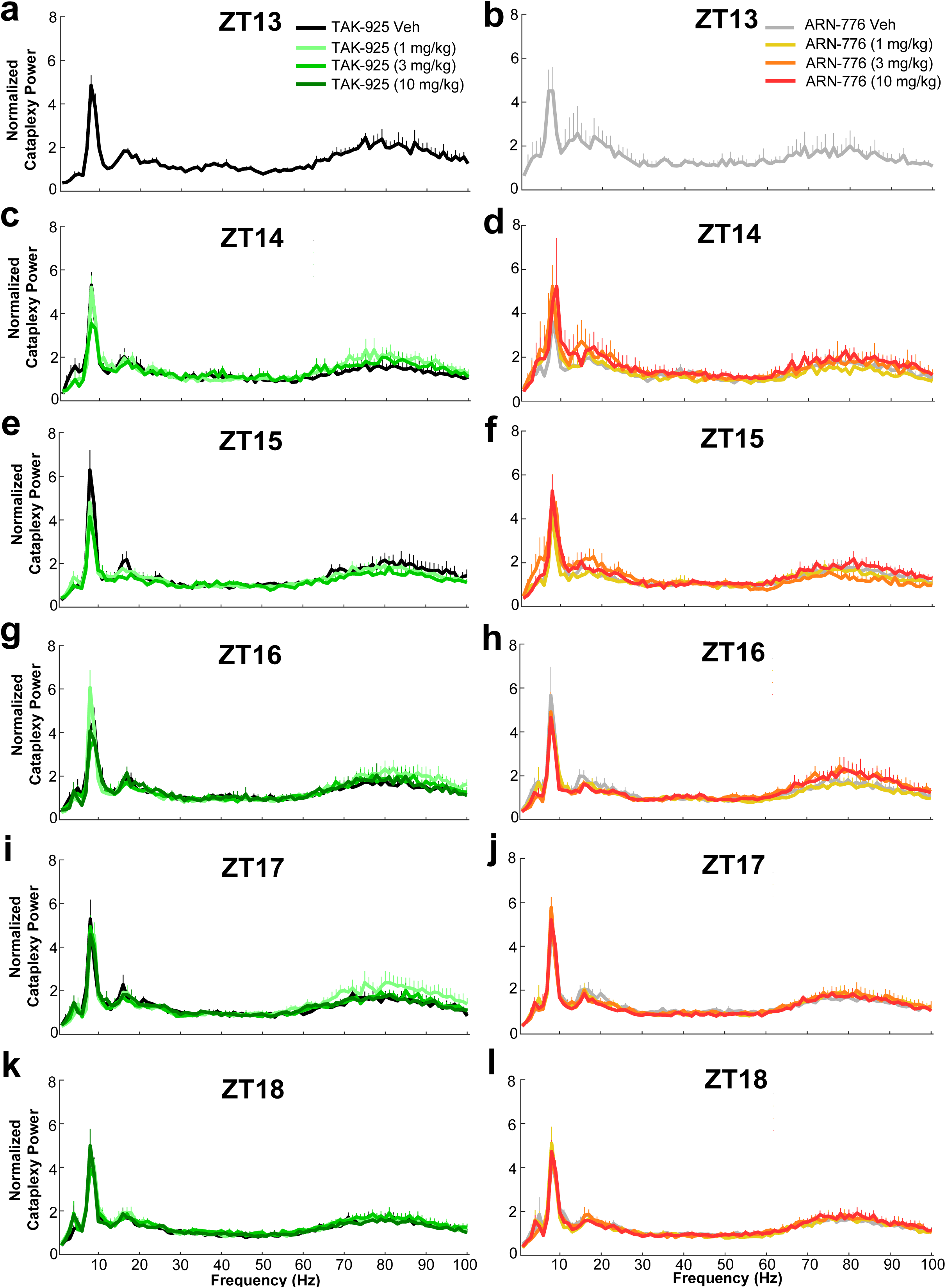
Normalized EEG spectral power (0 – 100 Hz) recorded during Cataplexy for the 6-h post-dosing recording period in male narcoleptic *orexin/tTA; TetO-DTA* mice treated with TAK-925 (left column) and ARN-776 (right column). Values are mean ± SEM. Abbreviations: Veh, vehicle; ZT, Zeitgeber Time; Hz, Hertz.

**Fig. 10.**
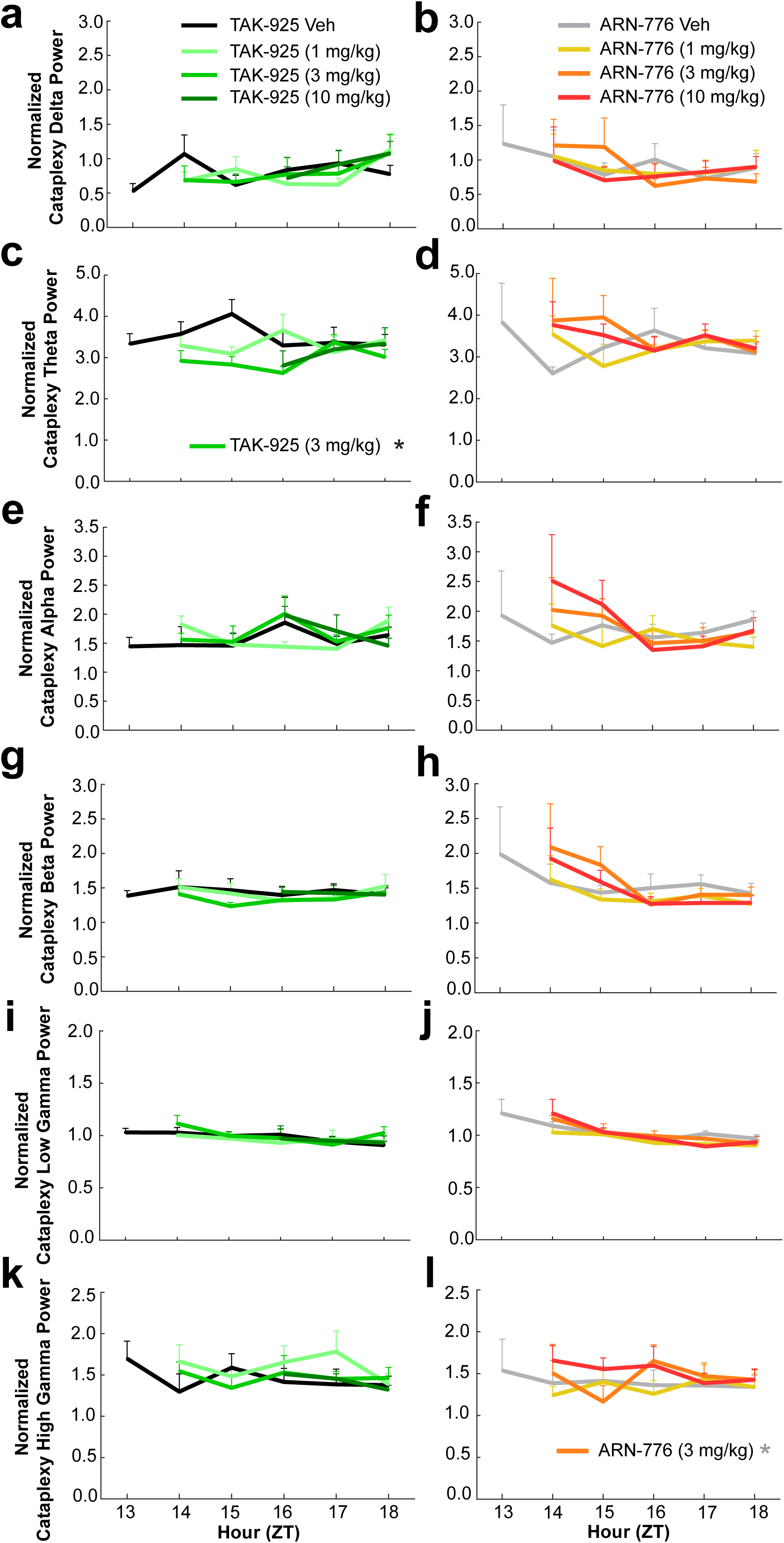
EEG spectra during Cataplexy for the conventional bandwidths in narcoleptic male *orexin/tTA; TetO-DTA* mice treated with TAK-925 and ARN-776**. a and b:** delta (0.5-4.0 Hz); **c and d:** theta (4-9 Hz); **e and f:** alpha (9-12 Hz); **g and h:** beta (12-30 Hz); **i and j:** low gamma (30-60 Hz); and **k and l:** high gamma (60-100 Hz). Values are mean ± SEM. * in legend denotes significance for that treatment across the 6-h recording as determined by 2-way ANOVA. Abbreviations: Veh, vehicle; ZT, Zeitgeber Time.

Due to the infrequent occurrences of Cataplexy during some hours, ANOVA did not calculate treatment x time effects.

### Activity Measures and T_sc_

TAK-925 and ARN-776 increased both gross motor activity (Fig. 11a and 11b) measured by the implanted telemeter and running wheel activity (Fig. 11c and 11d) during the first post-dosing hour. Two-way ANOVA indicated significant treatment x time effects on gross motor activity (*F*_(35, 210)_ *=* 2.49*, p =* 3.52 x 10^-5^); *post hoc* tests revealed significantly increased motor activity for all doses of TAK-925 as well as the 3 mg/kg and 10 mg/kg doses of ARN-776 during the first post-dosing hour (ZT13). Subsequent to this increase, reduced gross motor activity occurred later in the night for the highest doses of TAK-925 (ZT17; Fig. 11a) and ARN-776 (ZT15; Fig. 11b).

**Fig. 11.**
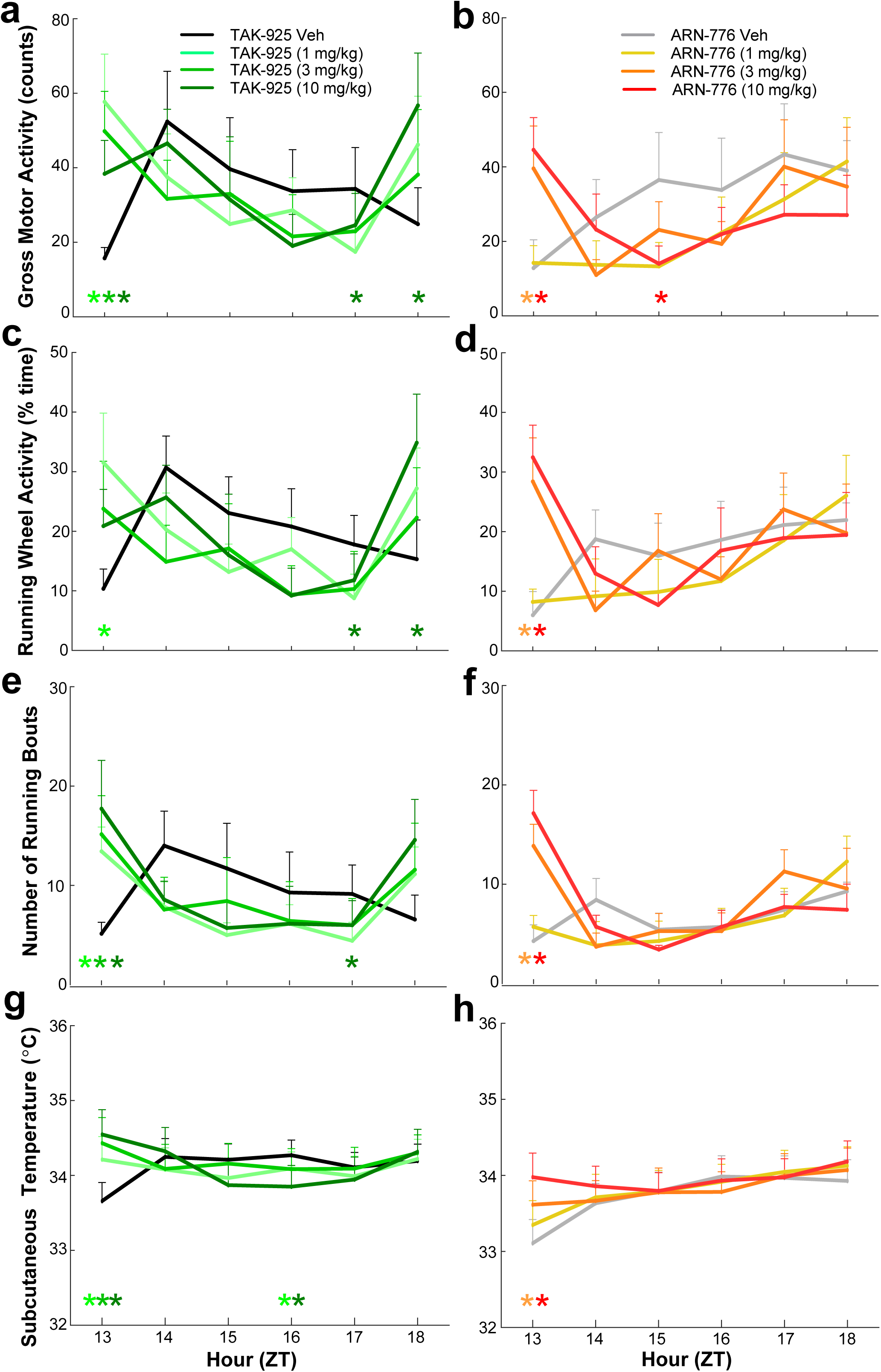
Gross motor activity (**a, b**), running wheel activity (**c, d**), number of running wheel bouts (**e, f**), and subcutaneous temperature (**g, h**) in narcoleptic male *orexin/tTA; TetO-DTA* mice treated with TAK-925 and ARN-776. Values are mean ± SEM. Colored * above hourly graphs indicate significance (*p*<0.05) during that hour relative to vehicle treatment. Abbreviations: Veh, vehicle; ZT, Zeitgeber Time.

Two-way ANOVA also indicated significant treatment x time effects on running wheel activity (*F*_(35, 210)_ *=* 2.33*, p =* 1.32 x 10^-4^); *post hoc* tests revealed significantly increased running wheel activity during the first post-dosing hour (ZT13) for the 1 mg/kg dose of TAK-925 (Fig. 11c) as well as for the 3 mg/kg and 10 mg/kg doses of ARN-776 (Fig. 11d). Importantly, the increased running during ZT13 was due to an increased number of running bouts (*F*_(35, 210)_ *=* 2.56*, p =* 2.02 x 10^-5^); *post hoc* tests confirmed more running bouts for all doses of TAK-925 (Fig. 11e) and for the 3 mg/kg and 10 mg/kg doses of ARN-776 (Fig. 11f). Reduced running wheel activity occurred later during ZT17 for TAK-925 (10 mg/kg) followed by an increase at ZT18.

The increases in activity early in the dark period were accompanied by elevated T_sc_ (Fig. 11g and 11h). Two-way ANOVA indicated both significant treatment (*F*_(7, 42)_ *=* 6.10*, p =* 5.97 x 10^-5^) and treatment x time (*F*_(35, 210)_ *=* 6.78*, p <* 3.34 x 10^-13^) effects on T_sc_*. Post hoc* tests revealed significantly increased T_sc_ for all doses of TAK-925 as well as for the 3 mg/kg and 10 mg/kg doses of ARN-776 during the first post-dosing hour (ZT13).

## Discussion

Current treatments for narcolepsy are directed toward symptomatic management of EDS and cataplexy. Amphetamines and other stimulants with abuse potential treat EDS through presynaptic stimulation of dopamine (DA) transmission [36, 39, 40]. Modafinil has fewer side effects than amphetamine-like stimulants and is the first-line treatment for EDS [36, 39, 40]. However, the norepinephrine-dopamine reuptake inhibitor solriamfetol and the histamine H3 antagonist/inverse agonist pitolisant have recently been approved by the FDA and EMEA to treat sleepiness associated with narcolepsy but do not appear to have stimulant effects [41]. Antidepressants, both tricyclic and selective monoaminergic reuptake inhibitors, alleviate cataplexy to the extent that they, or their metabolites, inhibit norepinephrine and serotonin uptake [36, 42–44]. Although gamma-hydroxybutyrate (GHB; aka sodium oxybate), a controlled substance with a difficult dosing regimen, was the only approved drug to treat both cataplexy and EDS for many years [45, 46], some of its limitations have been overcome by the combination of calcium oxybate/magnesium oxybate/potassium oxybate/sodium oxybate known as Xywav^→^ that was approved by the FDA in 2020.

In contrast to these symptomatic treatments, genetic and pharmacological rescue experiments [20–25] have established the proof-of-concept for Hcrt peptide replacement as the ideal therapy for narcolepsy. To date, however, only two small molecule HcrtR/OX2R agonists have been published, YNT-185 [29, 30] and TAK-925 [31]. In the present study, we evaluated the efficacy of TAK-925 and a related Hcrt agonist, ARN- 776.

### Physiological Efficacy Despite Limited Brain Penetration

Although our pharmacokinetic studies indicated that TAK-925 and ARN-776 showed limited penetration into the brain, both molecules produced clear physiological effects. Fujimoto and colleagues recently reported total plasma and brain levels of TAK-925 in C57BL/6 mice after intraperitoneal administration and found brain concentrations ranging from 183 to 63 ng/g over the first hour with a total brain:plasma concentration ratio of 0.2 based on data at two timepoints [35]. Despite differences in route of administration and mouse strain, our data are in general agreement with their findings. We observed total brain concentrations over the first hour after s.c. administration of TAK-925 to be 259 to 53 ng/g. Based on AUC values for free drug levels our free brain: plasma ratio is only 0.07:1, but this reflects a difference in free fraction for plasma versus brain as well as data over multiple timepoints. Similar to their findings, our data also demonstrated that TAK-925 as well as ARN-776 promoted wakefulness, but also increased activity and T_sc_ while suppressing NREM and REM sleep as well as cataplexy during the first hour after dosing. These results underscore that it is important to evaluate not only the brain to blood ratio of compounds but also their absolute free brain concentrations to determine the potential for target engagement.

### Similarities between TAK-925 and ARN-776

Although a direct “head to head” comparison of these two compounds was not possible due to differences in vehicle and route of administration, TAK-925 and ARN- 776 nonetheless produced remarkably similar results. As stated above, both compounds delayed sleep onset in a dose-related manner and, for at least the first hour after dosing, consolidated wakefulness into long wake bouts; reduced cataplexy, NREM and REM sleep; and increased activity, T_sc_ and spectral power in the alpha and gamma bands of the Waking EEG. All doses of TAK-925 and ARN-776 prolonged Wake Bout Duration across the entire 6-h recording. Furthermore, the reduction in cataplexy was sustained across the 6-h recording period at the highest dose of TAK-925 (10 mg/kg, s.c.) and all doses of ARN-776 (1-10 mg/kg, i.p.). Both compounds also had strong effects on the EEG during cataplexy. Lastly, despite the increased wakefulness during the first post-dosing hour, there was no compensatory increase in NREM sleep in the subsequent hours. As such, both of these putative Hcrt receptor agonists appear to have beneficial effects on wakefulness and cataplexy and thus encourage further development of Hcrt agonists as potential narcolepsy therapeutics.

### Differences between TAK-925 and ARN-776

Although it can be difficult to directly compare the efficacy of compounds, particularly since both the route of delivery (s.c. vs. i.p.) and the vehicles used to deliver these compounds differed, TAK-925 appeared to be more potent than ARN-776 on several measures. For example, although both compounds were tested at 1, 3 and 10 mg/kg, all doses of TAK-925 produced continuous wakefulness during the first hour after dosing whereas the wake-promoting effects of ARN-776 were dose-related.

Furthermore, much longer sleep onset latencies were observed with the lower doses of TAK-925 than after comparable doses of ARN-776. Although neither compound provoked an increase in NREM sleep subsequent to the increased wakefulness during the first dosing hour, the suppression of REM sleep by TAK-925 was more sustained and resulted in a significant REM rebound later in the night (Figs. 2g and 3e). Despite the absence of a NREM sleep rebound, the two compounds had very different effects on the EEG spectra during NREM sleep in the hour after the increased wakefulness (ZT14): whereas the 10 mg/kg dose of TAK-925 increased normalized EEG spectral power in all but the low gamma range, the highest dose of ARN-776 reduced normalized EEG spectral power in the alpha and high gamma range (Fig. 8). Although it is thus tempting to speculate that NREM sleep might thus be less restful after TAK- 925, EEG delta power, a frequency band associated with deep sleep, was increased during ZT14 after TAK-925. Although several of these physiological responses suggest a more prolonged effect of TAK-925, the different effects could be due to differential rates of clearance of these compounds due to the route and/or vehicles used for administration rather than to differences in the intrinsic activities of the two molecules.

### Perspective for Hcrt Agonist-based Therapeutics for Narcolepsy

Regarding potential clinical application, although both compounds effectively reduced cataplexy, the effects on wakefulness were primarily restricted to the first hour after dosing and may thus have limited utility to combat the EDS of narcolepsy patients. In addition, both agonists elevated gross motor and running wheel activity along with T_b,_ which does not occur with some wake-promoting agents that we have tested. The increased running wheel activity during the first hour after drug administration was due to an increased number of running bouts rather than fewer, prolonged bouts. These results suggest the possibility that the transient increase in wakefulness (and possibly alertness, as suggested by increased spectral power in the gamma bands of the EEG) was associated with elevated activity levels. Nonetheless, the anti-cataplectic activity of TAK-925 and ARN-776 is encouraging for the development of Hcrt receptor agonists to treat this pathognomonic symptom of narcolepsy and suggests that these particular molecules may be useful as tool compounds rather than as clinical development candidates. A Phase 1 study conducted in healthy sleep-deprived adults demonstrated that TAK-925 was well-tolerated and improved individuals’ ability to maintain wakefulness [51]. If these results were confirmed in a broader Phase 2 study and the drug was found to be safe and effective, TAK-925 could become the first narcoleptic therapeutic directed toward the neurotransmitter system whose dysfunction is implicated in the etiology of the disorder. However, TAK-925 requires intravenous administration, which has driven Takeda to pursue development of other agonists more suitable for clinical application.

Recently, Takeda has described an orally available HCRTR2/OX_2_R agonist, TAK-994, with an EC_50_ on OX_2_R = 19 nM and >700-fold selectivity against OX_1_R [52]. Oral administration of TAK-994 at 5h into the light period promoted wakefulness in wild type mice during their normal sleep phase [53]. Oral administration of TAK-994 during the active phase in both *orexin/ataxin-3* [16] and *orexin-tTA;TetO DTA* [19] narcolepsy mouse models significantly increased time spent in wake and improved wake maintenance while suppressing cataplectic episodes [52]. TAK-994 had wake-promoting effects following chronic dosing for up to 14 days in *orexin/ataxin-3* mice without causing a sleep rebound [54]. Unfortunately, a safety signal emerged in Phase 2 studies of TAK-994 which led Takeda to suspend dosing. Nevertheless, Takeda continues to advance other oral Hcrt receptor agonists, and other novel hypocretin agonists like ARN-776 continue are under development by other organizations as potential therapeutics for the treatment of narcolepsy, as well as for other disorders characterized by the presence of excessive daytime sleepiness.

## Acknowledgements

Research reported in this publication was supported by the National Institute of Neurological Disorders and Stroke of the National Institutes of Health R21 NS106882 and R01 NS098813 to T.S.K. The content is solely the responsibility of the authors and does not necessarily represent the official views of the National Institutes of Health. J.K.D.B. holds the Julie and Louis Beecherl, Jr., Chair in Medical Science and acknowledges support from the Robert A. Welch Foundation (grant I-1422). D.M.R. acknowledges support from the Robert A. Welch Foundation (grant I-1770). We thank Akihiro Yamanaka of Nagoya University for the C57BL/6-Tg(*orexin/tTA; TetO diphtheria toxin A* fragment)/Yamanaka mice and Haley Courtney, Laure Alexandre, Francisco Ortiz, and the institutionally-supported Preclinical Pharmacology Core at UT Southwestern Medical Center for technical assistance.

