## Supplementary Material for "Evaluation of the Efficacy of the Hypocretin/orexin Receptor Agonists TAK-925 and ARN-776 in Narcoleptic *Orexin/tTA; TetO-DTA* Mice"

**for**

**Synthesis of (2*R*,3*S*)-N-ethyl-2-((((1*s*,4*S*)-4-isopropylcyclohexyl)oxy)methyl)-3-(methylsulfonamido)piperidine-1-carboxamide (ARN-776)**

**3-bromo-2-((((1*r*,4*r*)-4-isopropylcyclohexyl)oxy)methyl)pyridine**

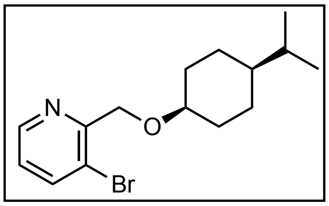

To a suspension of 60% sodium hydride (3.5 g, 87.5 mmol) in tetrahydrofuran (THF, 40 mL) was added cis-4-isopropylcyclohexanol (10.0 g, 70.3 mmol) at room temperature. The reaction mixture was stirred at room temperature for 12 h, after which 3-bromo-2-(bromomethyl)pyridine (8.7 g, 34.7 mmol) was added to the reaction mixture and the resulting suspension was stirred additional 12 h at room temperature. After completion, the reaction mixture was poured into a separatory funnel containing EtOAc (150 mL) and a 3:2:1 mixture of saturated aqueous $NH_4Cl$/brine/water (250 mL) (caution: vigorous bubbling ensues upon addition). The layers were separated and the aqueous layer was extracted with EtOAc (3x200 mL). The combined organic layers were washed with water (100 mL), brine (100 mL), dried over $Na_2SO_4$, filtered, and concentrated to afford crude product. Purification was accomplished by silica gel flash column chromatography (hexanes:EtOAc = 9:1) affording the title compound, 3-bromo-2-((((1*r*,4*r*)-4-isopropylcyclohexyl)oxy)methyl)pyridine (8.6 g, 80%) as a viscous oil. Compound purity was established by TLC (one spot) analysis.

R*f*: 0.50 (hexane:EtOAc = 9:1); $^1$H NMR (400 MHz, $CDCl_3$) δ 8.45 (dd, *J* = 4.7, 1.5 Hz, 1H), 7.77 (dd, *J* = 8.0, 1.5 Hz, 1H), 7.02 (dd, *J* = 8.0, 4.7 Hz, 1H), 4.62 (s, 2H), 3.65 (dq, *J* = 5.1, 2.6 Hz, 1H), 1.86 – 1.98 (m, 2H), 1.26 – 1.51 (m, 7H), 0.96 – 1.06 (m, 1H), 0.79 (d, *J* = 6.9 Hz, 6H); $^{13}$C NMR (101 MHz, $CDCl_3$) δ 156.3, 147.5, 140.4, 123.8, 121.5, 74.5, 70.8, 43.2, 31.9, 29.7, 23.9, 19.8; MS(ES) calculated for $C_{15}H_{22}BrNO$ $[M + H]^+$ 312.10; found 312.2, 314.20.

PROTON_01

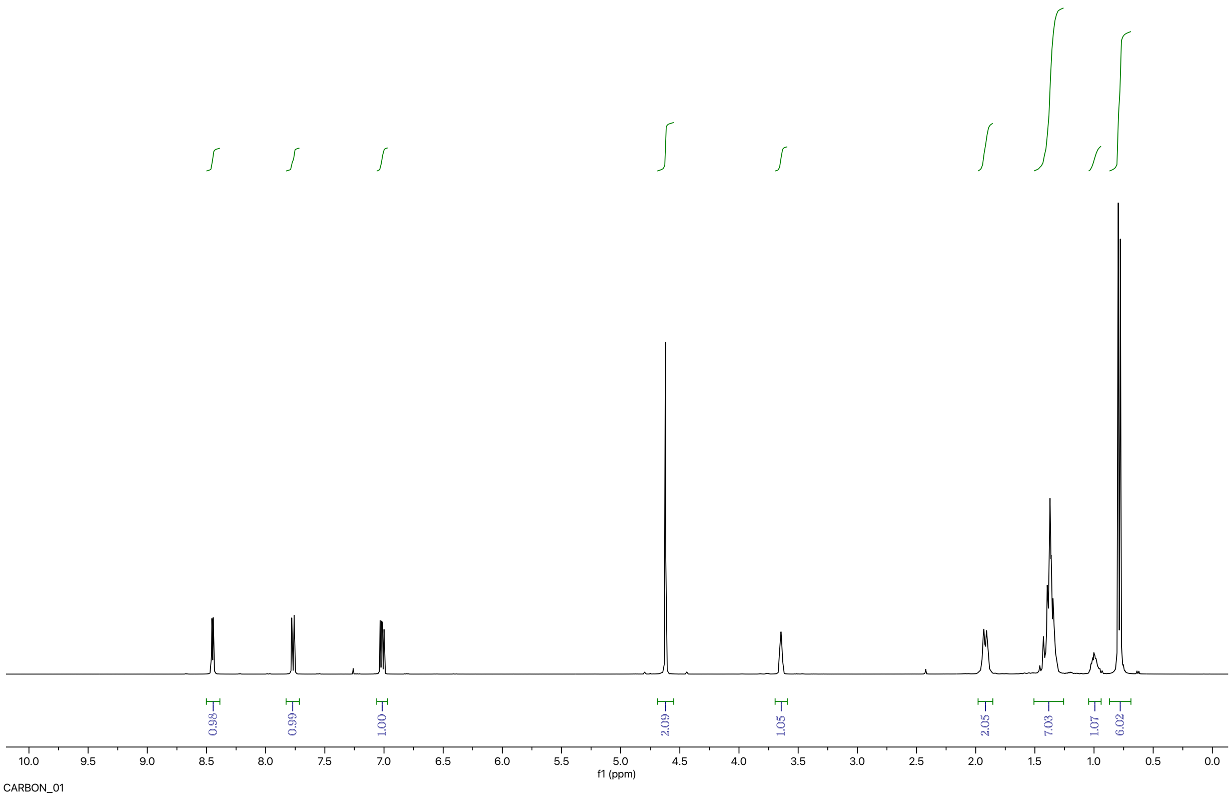


CARBON_01

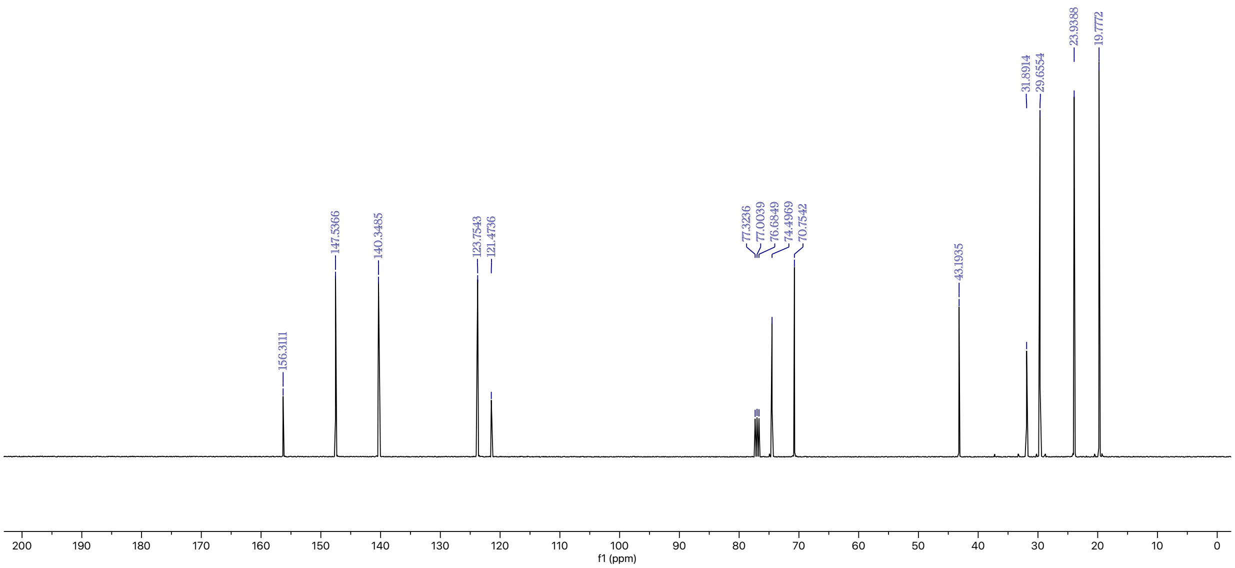

***N*-(2-((((1*s*,4*s*)-4-isopropylcyclohexyl)oxy)methyl)pyridin-3-yl)methanesulfonamide**

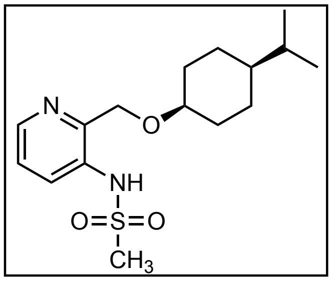


To a flame-dried, round-bottom flask equipped with reflux condenser and magnetic stir bar was sequentially added 3-bromo-2-(((cis-4-isopropylcyclohexyl)oxy)methyl)pyridine (3.0 g, 9.6 mmol), methanesulfonamide (1.1 g, 11.6 mmol), di-tertbutyl(2′,4′,6′-triiso propylbiphenyl2-yl)phosphine (0.41 g, 1.0 mmol), tris(dibenzylidene acetone)dipalladium(0) (0.44 g, 0.5 mmol), cesium carbonate (4.7 g, 14.4 mmol) and THF (40 mL). The reaction mixture was refluxed at 80 °C for 12 h, upon which TLC analysis indicated complete conversion of starting material to product. The reaction mixture was filtered through celite, and the filtrate was extracted with EtOAc. The reaction mixture was poured into a separatory funnel containing saturated aqueous brine (200 mL). The layers were separated, and the aqueous layer was extracted with EtOAc (3x200 mL). The combined organic layers were dried over $Na_2SO_4$, filtered, and concentrated to afford a crude product. Purification was accomplished by silica gel flash column chromatography (hexanes:EtOAc = 1:1) affording *N*-(2-((((1*s*,4*s*)-4-isopropylcyclohexyl)oxy)methyl)pyridin-3-yl)methanesulfonamide (2.3 g, 74%) as an amorphous solid. Compound purity was established by TLC (one spot) analysis. R*f*: 0.40 (hexanes:EtOAc = 1:1); $^1$H NMR (400 MHz, $CDCl_3$) δ 8.70 (s, 1H), 8.20 (dd, *J* = 4.7, 1.4 Hz, 1H), 7.84 (dd, *J* = 8.3, 1.5 Hz, 1H), 7.17 (dd, *J* = 8.3, 4.7 Hz, 1H), 4.73 (s, 2H), 3.56 – 3.69 (m, 1H), 2.96 (s, 3H), 1.91 (brd, *J* = 12.8 Hz, 2H), 1.33 – 1.48 (m, 5H), 1.19 – 1.30 (m, 2H), 0.93 – 1.05 (m, 1H), 0.78 (d, *J* = 6.9 Hz, 6H); $^{13}$C NMR (101 MHz, $CDCl_3$) δ 146.6, 144.0, 133.9, 126.8, 123.2, 75.2, 71.8, 43.1, 39.9, 32.0, 29.3, 23.8, 19.7; MS(ES) calculated for $C_{16}H_{26}N_2O_3S$ $[M + H]^+$ 327.17, found 327.30

PROTON_01

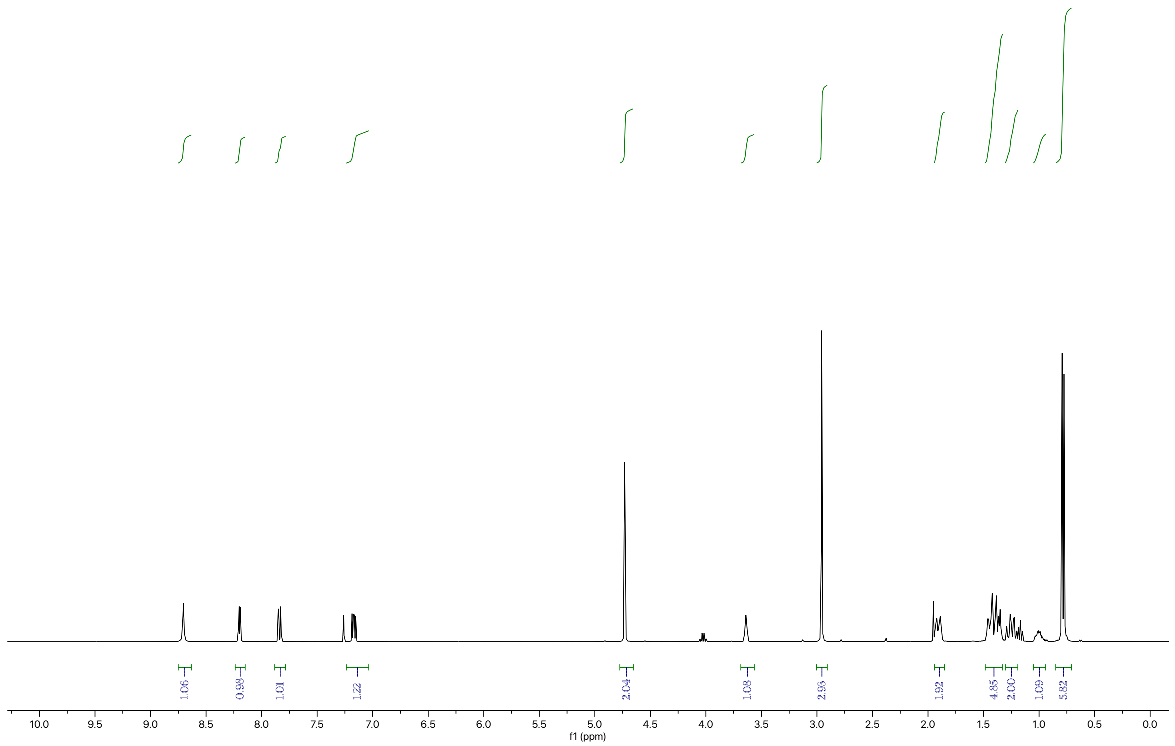


CARBON_01

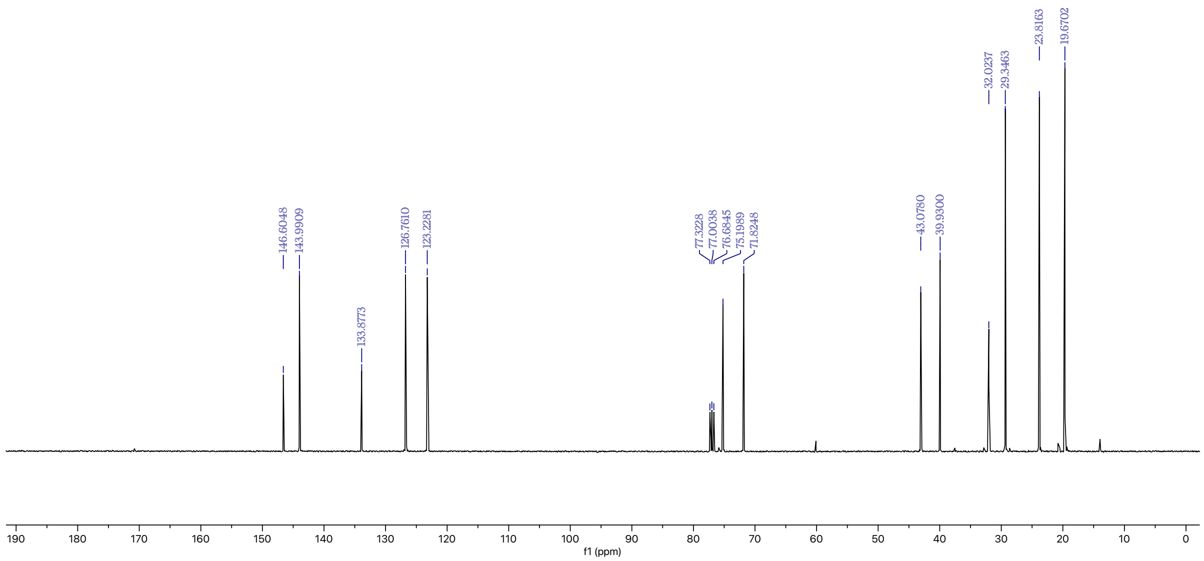

***N*-(2-((((1*s*,4*s*)-4-isopropylcyclohexyl)oxy)methyl)piperidin-3-yl)methanesulfonamide**

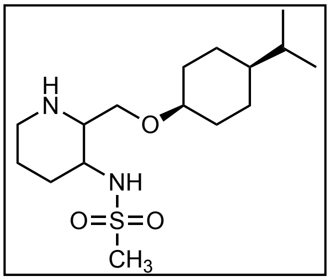

(2-(((cis-4-isopropylcyclohexyl)oxy)methyl)pyridin-3-yl)methane sulfonamide (2.3 g, 7.0 mmol, platinum oxide (0.08 g, 0.35 mmol), methanol (15.0 mL) and acetic acid (15.0 mL) was stirred overnight under a 0.6 MPa hydrogen atmosphere at 50 °C. The mixture was filtrated, and the filtrate was neutralized with saturated aqueous sodium hydrogen carbonate ($NaHCO_3$) solution at 0 °C and extracted with EtOAc (3x100 mL). The combined organic layers washed with water, saturated brine, dried over $Na_2SO_4$, filtered, and concentrated under reduced pressure. Purification was accomplished by silica gel flash column chromatography (hexanes:EtOAc = 1:1) affording the title compound (1.6 g, 70%) as an amorphous solid. Compound purity was established by TLC (one spot) analysis.

R*f*: 0.35 (hexanes:EtOAc = 1:1); $^1$H NMR (400 MHz, $CDCl_3$) δ 3.61 (dd, *J* = 9.7, 3.7 Hz, 1H), 3.53 (dd, *J* = 9.7, 5.7 Hz, 1H), 3.48 (brs, 1H), 3.34 (td, *J* = 9.6, 4.0 Hz, 1H), 3.06 (dt, *J* = 12.5, 3.9 Hz, 1H), 2.92 (s, 3H), 2.79 (ddd, *J* = 9.3, 5.6, 3.8 Hz, 1H), 2.64 (td, *J* = 12.2, 11.8, 3.1 Hz, 1H), 2.09 – 2.20 (m, 1H), 1.92 (s, 2H, $CH_3CO_2H$ residue), 1.70 – 1.86 (m, 3H), 1.52 – 1.65 (m, 1H), 1.16 - 1.51 (m, 8H), 0.91 – 1.04 (m, 1H), 0.79 (d, *J* = 6.8 Hz, 6H); $^{13}$C NMR (101 MHz, $CDCl_3$) δ 74.9, 66.6, 59.2, 51.0, 43.6, 43.1, 41.3, 32.0, 31.9, 29.5, 24.0, 23.6, 22.6, 19.8($CH_3CO_2H$ residue); MS(ES) calculated for $C_{16}H_{32}N_2O_3S$ $[M + H]^+$ 333.22, found 333.20.

PROTON_01

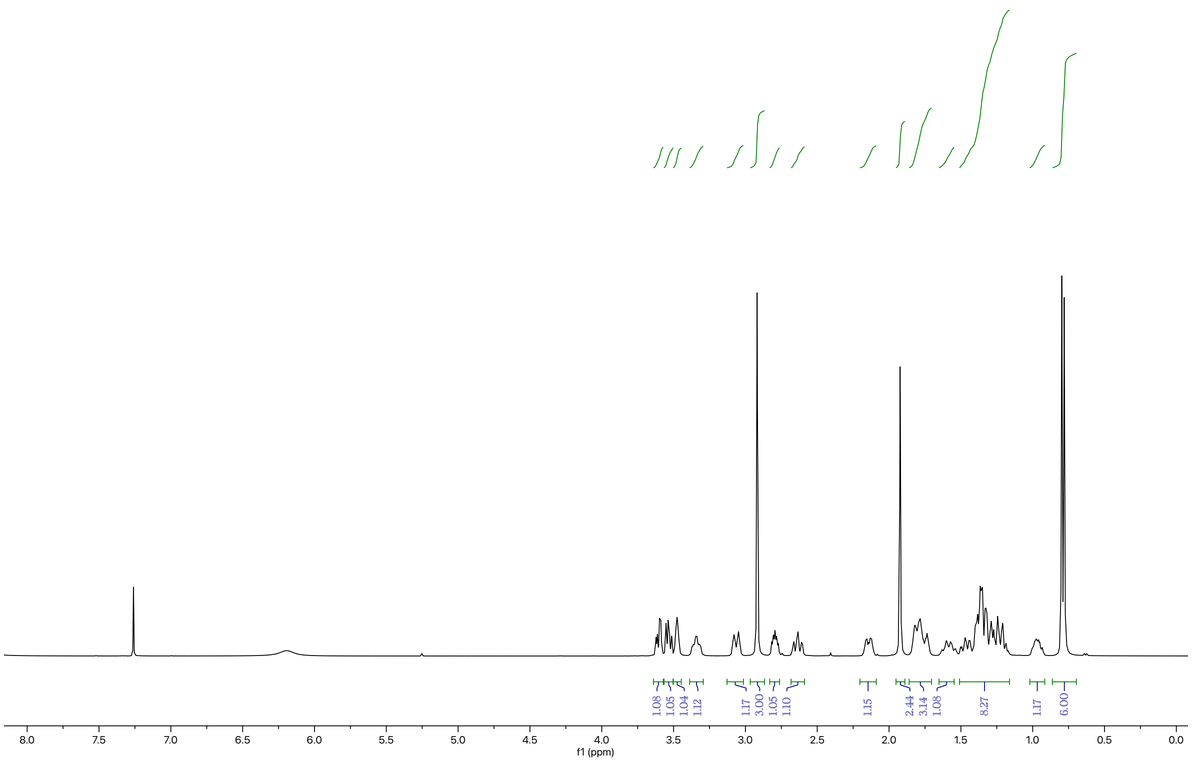


CARBON_01

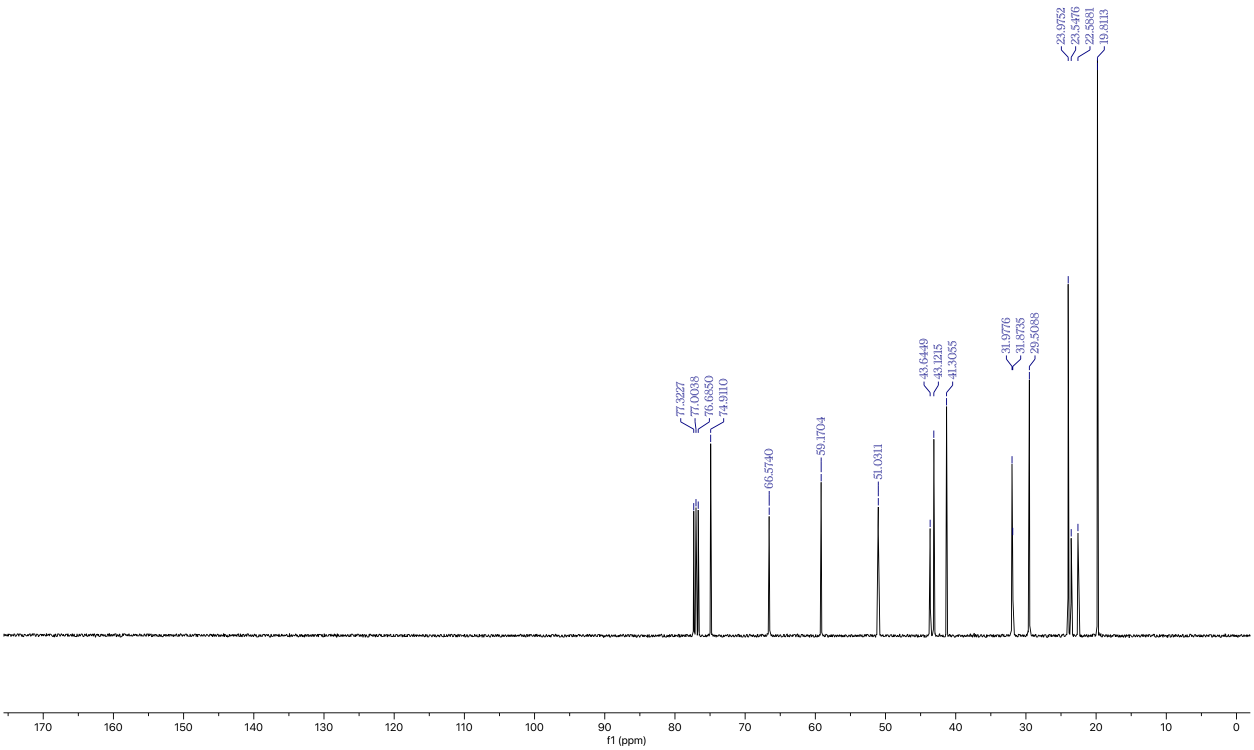

**(2*R*,3*S*)-*N*-ethyl-2-((((1*s*,4*S*)-4-isopropylcyclohexyl)oxy)methyl)-3-(methylsulfonamido)piperidine-1-carboxamide**

To a solution of (*2S,3S*)-2,3-bis((4-methylbenzoyl)oxy)succinic acid (579 mg, 1.5 mmol) in

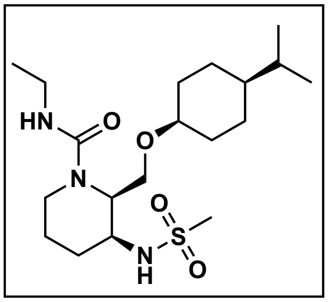


ethanol (4 ml) was added a solution of *N*-(cis-2-(((cis-4-isopropyl cyclohexyl)oxy)methyl)piperidin-3-yl)methanesulfonamide (498 mg, 1.5 mmol) in ethanol (4 ml) at room temperature, and the solution was left standing overnight. The resulting solid was collected by filtration and washed with acetonitrile to give a solid (270 mg, 0.8 mmol) methanesulfonamide derivatives. To a solution of *N*-(cis-2-(((cis-4-isopropylcyclohexyl)oxy)methyl)piperidin-3-yl)methanesulfonamide (100.0 mg, 0.30 mmol) and $Et_3N$ (0.08 mL, 0.60 mmol) in THF (2.0 mL) was added ethyl isocyanate (21.3 mg, 0.30 mmol) at 0 °C, and the whole reaction mixture was warmed it to room temperature and stirred for 12 h. After completion, the reaction mixture was quenched with the addition of water (10 mL) and extracted with ethyl acetate (3x50 mL). The organic layer was washed with saturated brine, dried over anhydrous sodium sulfate ($Na_2SO_4$), and the solvent was evaporated under reduced pressure. The residue was purified by silica gel flash chromatography (hexanes:EtOAc = 2:3) to give the title compound (2*R*,3*S*)-*N*-ethyl-2-((((1*s*,4*S*)-4-isopropylcyclohexyl)oxy)methyl)-3-(methylsulfonamido)piperidine-1-carboxamide (60.0 mg, 50% yield) as solid.

R*f*: 0.35 (hexanes:EtOAc = 2:3); $^1H$ NMR (400 MHz, $CDCl_3$) δ 5.83 (d, *J* = 7.2 Hz, 1H), 4.75 – 4.87 (m, 1H), 4.43 – 4.56 (m, 1H), 3.87 (dd, *J* = 9.2, 7.5 Hz, 1H), 3.66 - 3.80 (m, 1H), 3.47 – 3.59 (m, 3H), 3.17 – 3.32 (m, 2H), 3.00 (s, 3H), 2.74 – 2.86 (m, 1H), 1.84 – 1.91 (m, 2H), 1.56 – 1.80 (m, 3H), 1.34 – 1.52 (m, 5H), 1.10 – 1.28 (m, 5H), 0.99 – 1.08 (m, 1H), 0.86 (d, *J* = 6.8 Hz, 6H); $^{13}C$ NMR (101 MHz, $CDCl_3$) δ 158.1, 75.1, 64.4, 53.3, 52.6, 43.2, 40.9, 38.7, 35.7, 32.2, 29.6, 29.4, 27.9, 24.6, 23.9, 23.8, 19.7, 19.7, 15.2; MS(ES) calculated for $C_{19}H_{37}N_3O_4S$ $[M + H]^+$ 404.26, found 404.20.

*Ref.* Takeda Pharmaceutical Company Limited; WO2017135306 - Substituted piperidine compound and use thereof

ARN-1-776-2
STANDARD FLUORINE PARAMETERS

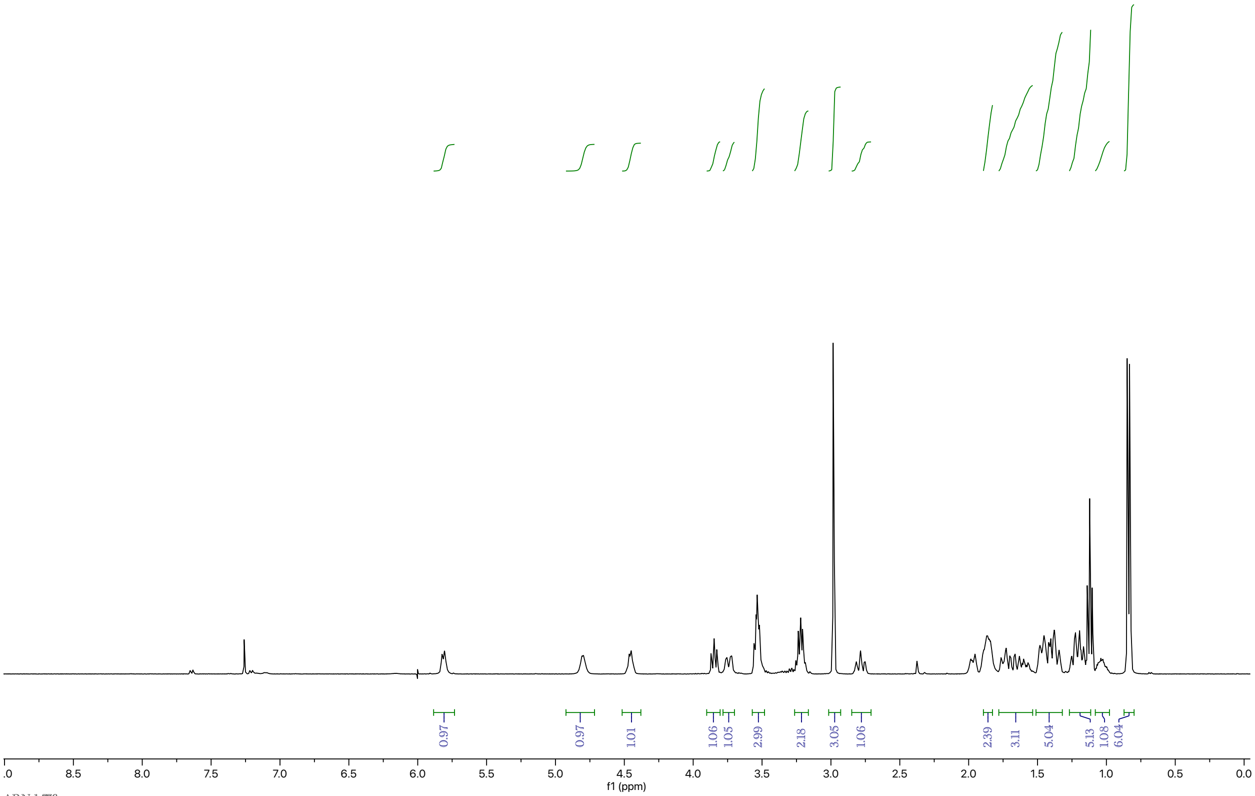


ARN-1-776c

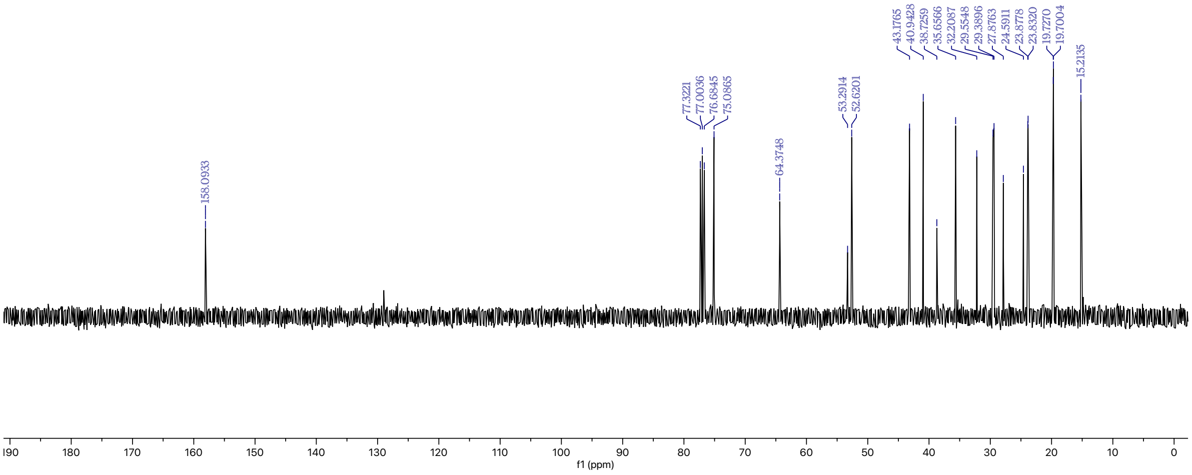
